# Bistability and oscillations in cooperative microtubule and kinetochore dynamics in the mitotic spindle

**DOI:** 10.1101/575530

**Authors:** Felix Schwietert, Jan Kierfeld

## Abstract

In the mitotic spindle microtubules attach to kinetochores via catch bonds during metaphase. We investigate the cooperative stochastic microtubule dynamics in spindle models consisting of ensembles of parallel microtubules, which attach to a kinetochore via elastic linkers. We include the dynamic instability of microtubules and forces on microtubules and kinetochores from elastic linkers. We start with a one-sided model, where an external force acts on the kinetochore. A mean-field approach based on Fokker-Planck equations enables us to analytically solve the one-sided spindle model, which establishes a bistable force-velocity relation of the microtubule ensemble. All results are in agreement with stochastic simulations. We derive constraints on linker stiffness and microtubule number for bistability. The bistable force-velocity relation of the one-sided spindle model gives rise to oscillations in the two-sided model, which can explain stochastic chromosome oscillations in metaphase (directional instability). We also derive constraints on linker stiffness and microtubule number for metaphase chromosome oscillations. We can include poleward microtubule flux and polar ejection forces into the model and provide an explanation for the experimentally observed suppression of chromosome oscillations in cells with high poleward flux velocities. Chromosome oscillations persist in the presence of polar ejection forces, however, with a reduced amplitude and a phase shift between sister kinetochores. Moreover, polar ejection forces are necessary to align the chromosomes at the spindle equator and stabilize an alternating oscillation pattern of the two kinetochores. Finally, we modify the model such that microtubules can only exert tensile forces on the kinetochore resulting in a tug-of-war between the two microtubule ensembles. Then, induced microtubule catastrophes after reaching the kinetochore are necessary to stimulate oscillations.

**Author summary:** The mitotic spindle is responsible for proper separation of chromosomes during cell division. Microtubules are dynamic protein filaments that actively pull chromosomes apart during separation. Two ensembles of microtubules grow from the two spindle poles towards the chromosomes, attach on opposite sides, and pull chromosomes by depolymerization forces. In order to exert pulling forces, microtubules attach to chromosomes at protein complexes called kinetochores. Before the final separation, stochastic oscillations of chromosomes are observed, where the two opposing ensembles of microtubules move chromosome pairs back an forth in a tug-of-war.

Using a a combined computational and theoretical approach we quantitatively analyze the emerging chromosome dynamics starting from the stochastic growth dynamics of individual microtubules. Each of the opposing microtubule ensembles is a bistable system, and coupling two such systems in a tug-of-war results in stochastic oscillations. We can quantify constraints on the microtubule-kinetochore linker stiffness and the microtubule number both for bistability of the one-sided system and for oscillations in the full two-sided spindle system, which can rationalize several experimental observations. Our model can provide additional information on the microtubule-kinetochore linkers whose molecular nature is not completely known up to now.

## 1 Introduction

Proper separation of chromosomes during mitosis is essential for the maintenance of life and achieved by the mitotic spindle, which is composed of two microtubule (MT) asters anchored at the spindle poles. The spindle contains three types of MTs classified according to their function [1]: astral MTs interact with the cell membrane to position the spindle poles, interpolar MTs interact with MTs from the opposite pole to maintain spindle length, and, finally, kinetochore MTs link to the chromosomes via the kinetochores at the centromere and can apply pulling forces via the linkage. The MT-kinetochore bond is a catch bond [2], i.e., tightening under tension but the molecular nature of the MT-kinetochore link is not exactly known and a complete mechanistic understanding of the catch bond is missing [3, 4] but probably involves Aurora B [5]; the Ndc80 complexes and Dam1 (in yeast) are believed to play a major role in the MT-kinetochore link. One function of the spindle is to align the chromosomes in the metaphase plate at the spindle equator. It has been observed in several vertebrate cells that chromosomes do not rest during metaphase but exhibit oscillations along the pole to pole axis known as directional instability [6–12], whereas in Drosophila embryos and Xenopus eggs a directional instability does not occur [13, 14]. If present, these oscillations are stochastic and on the time scale of minutes, i.e., on a much larger time scale than the dynamic instability of single MTs. Both single kinetochores and the inter-kinetochore distance oscillate; inter-kinetochore or breathing oscillations occur with twice the frequency of single kinetochore oscillations [11].

A quantitative understanding of mitosis and the underlying mechanics of the MT-kinetochore-chromosome system is still lacking. In the past, several theoretical models have been proposed (reviewed in [15, 16]). Many models that account for force balance in metaphase and directional instability have in common that they are one-dimensional, neglect spindle pole dynamics and are driven by dynamic instability [17] of MTs. Differences lie in the modeling of the MT-kinetochore link and the question how depolymerizing MTs can exert forces on the kinetochore. An early model by Joglekar and Hunt [18] uses the thermodynamic Hill sleeve model [19] for MT-kinetochore connection and reproduces directional instability. Later, Civelekoglu-Scholey *et al.* [20] proposed a model in which force is transmitted by motor proteins. By variation of the model parameters they were able to reproduce a wide range of chromosome behavior observed in different cell types. More recent studies, which have suggested Ndc80 fibrils as main force transmitter [4, 21, 22], motivated models in which the MTs are linked to the kinetochore via (visco-)elastic springs [23, 24]. A new model by Civelekoglu-Scholey *et al.* [23] uses viscoelastic bonds and accounts for the observation that in PtK1 cells only chromosomes in the center of the metaphase plate exhibit directional instability [11]. They explain this dichotomy with different distributions of polar ejection forces at the center and the periphery of the metaphase plate. Polar ejection forces originate from non-kinetochore MTs interacting with the chromosome arms and pushing them thereby towards the spindle equator. Banigan *et al.* [24] developed a minimal model with simple elastic linkers and neglecting polar ejection forces. Referring to experiments with budding yeast kinetochores [2], they modeled MT dynamics with force-dependent velocities, catastrophe and rescue rates. In simulations of one side of their model with one spindle pole and one kinetochore, Banigan *et al.* found a bistable behavior of the kinetochore velocity in dependence of an external force applied to the kinetochore. From this bistability, kinetochore oscillations in the full model can be deduced as the kinetochores periodically pass hysteresis loops. So far, stochastic chromosome oscillations have been reproduced in a number of models in simulations [18, 20, 23, 24] but a predictive theory, which establishes the connection between the dynamic instability of MTs and MT-kinetochore linker properties on the one hand and the stochastic oscillations on the other hand, is missing.

Here, we first derive an analytical solution of the one-sided model of Banigan *et al.* from a novel mean-field approach. For this purpose, we start from the Fokker-Planck equations for the length distribution of the MT-kinetochore linkers. The only mean-field approximation is to neglect stochastic velocity fluctuations of the attached kinetochore. The stationary state solution allows us to quantify for which model parameters a bistability in the force-velocity relation occurs in the one-sided model. In particular, we can quantify the bistable regime in the parameter plane of MT-kinetochore linker stiffness and MT numbers. Interpreting the force-velocity relation as phase space diagram for the two-sided model, we are able (1) to quantify an oscillatory regime, in which kinetochores exhibit directional instability, in the parameter plane of linker stiffness and MT numbers predicting that linkers have to be sufficiently stiff; (2) to describe kinetochore motion in this oscillatory regime, calculate frequencies which agree with in vivo measurements [11] and to explain frequency doubling of breathing compared to single kinetochore oscillations; (3) to describe kinetochore motion in the non-oscillatory regime as fluctuations around a fixed point; (4) to show that high poleward flux velocities move the system out of the oscillatory regime and thereby explain why directional instability has been observed in mitotic vertebrate cells but not in Drosophila embryos and Xenopus eggs; (5) to show that polar ejection forces reduce the amplitude of oscillations, induce a phase shift between sister kinetochores and are necessary to align the chromosome at the spindle equator; (6) to derive as necessary condition for oscillations that either MTs must be able to apply pushing forces on the kinetochore or a catastrophe has to be induced with increased catastrophe rate when a MT reaches the kinetochore. All these results are validated by stochastic simulations.

In particular, we quantify in (1) lower bounds for linker stiffnesses that allow oscillations, whose value depends on the behavior of MTs growing against the kinetochore. If MTs can exert pushing forces, we find oscillations for linker stiffnesses > 16 pN μm^−1^; also if MT catastrophes are induced upon reaching the kinetochore, we find oscillations in a similar range of linker stiffnesses. These constraints provide useful additional information on MT-kinetochore linkers whose molecular nature is not completely unraveled up to now.

## 2 Mitotic spindle model

We use a one-dimensional model of the mitotic spindle (Fig. 1A), similar to the model from Ref. [24]. The *x*-coordinate specifies positions along the one-dimensional model, and we choose *x* = 0 to be the spindle equator. The spindle model contains a single chromosome represented by two kinetochores, which are linked by a spring with stiffness *c*_k_ and rest length *d*_0_. Two centrosomes margin the spindle at ±*x*_c_. From each centrosome a constant number *M* of MTs emerges with their plus ends directed towards the spindle equator. Each MT exhibits dynamic instability [17] and attaches to and detaches from the corresponding kinetochore stochastically. Attached MTs are connected to the kinetochore by a linker, which we model as Hookean polymeric spring with stiffness *c* and zero rest length. This spring exerts a force *F*_mk_ = −*c*(*x*_m_ − *X*_k_) on each MT, and each MT exerts a counter force −*F*_mk_ on the kinetochore, where *X*_k_ and *x*_m_ are kinetochore and MT position.

**Fig 1.**
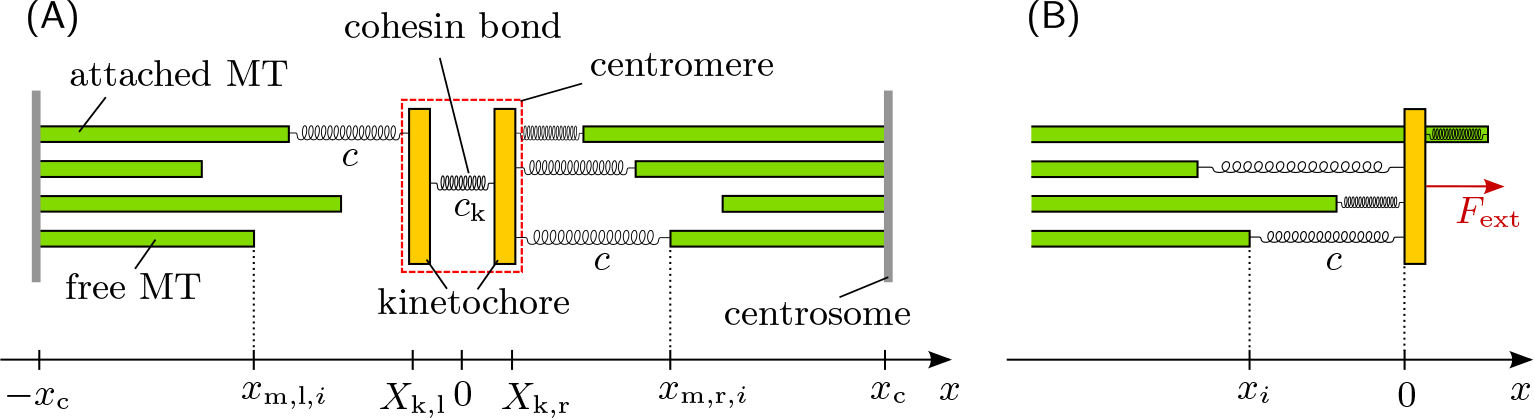
One-dimensional model of the mitotic spindle. (A) Two-sided model: *M* MTs arise from each centrosome and can attach to / detach from the corresponding kinetochore. (B) One-sided model: Left half of two-sided model. The cohesin bond is replaced by the external force *F*_ext_. MTs are not confined by a centrosome and permanently attached to the kinetochore. MT-kinetochore distances *x_i_* = *x*_m,*i*_ − *X*_k_ are the only relevant coordinates.

In the following we define all MT parameters for MTs in the left half of the spindle model; for MTs in the right half position velocities *v* and forces *F* have opposite signs. In the left half, tensile forces on the MT-kinetochore link arise for *X*_k_ > *x*_m_ and pull the MT in the positive *x*-direction, *F*_mk_ > 0. In Ref. [2], the velocities of MT growth *v*_m+_ and shrinkage *v*_m−_ as well as the rates of catastrophe *ω*_c_, rescue *ω*_r_ and detachment *ω*_d±_ have been measured while MTs were attached to reconstituted yeast kinetochores. They can all be described by an exponential dependence on the force *F*_mk_ that acts on the MT plus end:

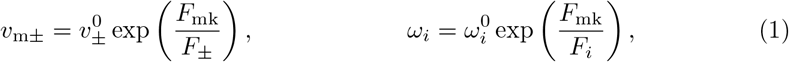

(for *i* = r, c, d+, d−) with *F*_+_, *F*_r_, *F*_d+_ > 0 and *F*_−_, *F*_c_, *F*_d−_ < 0 for the characteristic forces, because tension (*F*_mk_ > 0) enhances growth velocity, rescue and detachment of a growing MT, while it suppresses shrinking velocity, catastrophe and detachment of a shrinking MT (note that we use signed velocities throughout the paper, i.e., *v*_m−_ < 0 and *v*_m+_ > 0). Suppression of detachment is the catch bond property of the MT-kinetochore link. The attachment rate is assumed to follow a Boltzmann distribution,

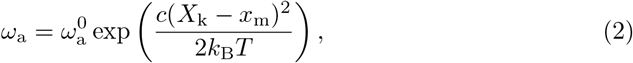

according to the MT-kinetochore linker spring energy.

The kinetochore motion is described by an overdamped dynamics,

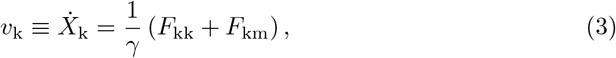

with the friction coefficient *γ*, and the forces *F*_kk_ and *F*_km_ = ∑_att. MTs_ *F*_mk_ originating from the cohesin bond between kinetochores and the MT-kinetochore linkers of all attached MTs, respectively. All experimental results for parameter values are listed in Table 1.

**Table 1.** Model parameters.

| **Transition parameters** | $\omega_i$ | $\omega_i^0$ in $\text{s}^{-1}$ | $F_i$ in pN |
| --- | --- | --- | --- |
| catastrophe | $\omega_c$ | 0.0019 | -2.3 [2] |
| rescue | $\omega_r$ | 0.024 | 6.4 [2] |
| detachment in growing state | $\omega_{d+}$ | 0.00011 | 3.8 [2] |
| detachment in shrinking state | $\omega_{d-}$ | 0.035 | -4.0 [2] |
| attachment rate | $\omega_a$ | 1.0 | estimated |
| **Velocity parameters** | $v_{m\pm}$ | $v_{\pm}^0$ in $\text{nms}^{-1}$ | $F_{\pm}$ in pN |
| growth | $v_{m+}$ | 5.2 | 8.7 [2] |
| shrinking | $v_{m-}$ | -210.0 | -3.2 [2] |
| **Other parameters** | Symbol | Value |  |
| cohesin bond stiffness | $c_k$ | 20 pN $\mu\text{m}^{-1}$ | estimated |
| cohesin bond rest length | $d_0$ | 1 $\mu\text{m}$ | [7] |
| centrosome position | $x_c$ | 6.8 $\mu\text{m}$ | [10] |
| friction coefficient | $\gamma$ | 1 pN s $\mu\text{m}^{-1}$ | estimated |
| thermal energy | $k_B T$ | 4 pN nm | estimated |

In the simple spring model for the MT-kinetochore linker the MT plus ends are able to “overtake” the kinetochore (*x*_m_ > *X*_k_, again for MTs in the left half of the spindle) and thereby exert pushing forces *F*_km_ > 0 on the kinetochore (which could be interpreted as a compression of the MT-kinetochore linker). It is questionable, however, whether kinetochore attachments are really able to produce such pushing forces [7, 25]. Therefore, we will later modify the model such that the growth of a MT is stalled or that the MT undergoes a catastrophe when it reaches the kinetochore. There are other forces that push the kinetochore away from the pole, e.g., the polar ejection forces transmitted by non-kinetochore MTs through collisions with the chromosome arms or via chromokinesins belonging to the kinesin-4 and kinesin-10 families [26]. We first study the model in the absence of polar ejection forces and include them in a second step into the model as external forces, which depend on the absolute positions of the kinetochores. We also neglect poleward microtubule flux in the beginning, which describes a constant flux of tubulin from the plus-ends towards the spindle pole and is probably driven by plus-end directed kinesin-5 motors pushing overlapping antiparallel MTs apart and kinesin-13 proteins that depolymerize MTs at the centrosome [27]. Poleward microtubule flux has a pulling effect on MTs and kinetochores and will also be included later on into our model by shifting the MT velocities *v*_m±_.

At the centrosome, MTs are confined: It is reasonable to assume that they undergo a forced rescue and detach from the kinetochore if they shrink to zero length. If the mean distance of MTs from the spindle equator is sufficiently small, |〈*x*_m_〉| ≪ |*x*_c_|, we can also consider the MTs as unconfined (|*x*_c_| → ∞). Then both MT and kinetochore dynamics solely depend on their relative distances and not on absolute positions, which simplifies the analysis.

## 3 Mean-field theory for bistability in the one-sided model

We first examine the one-sided model of Banigan *et al.* [24], which only consists of the left half of the two-sided model with an external force applied to the kinetochore (Fig. 1B). In simulations of this one-sided spindle model, kinetochore movement exhibits bistable behavior as a function of the applied force. This bistability can explain stochastic chromosome oscillations in metaphase. In the following, we present a Fokker-Planck mean-field approach that lets us derive bistability analytically and obtain constraints for the occurrence of bistability. We obtain a system of Fokker-Planck equations (FPEs) for the *M* MT-kinetochore distances *x_i_* ≡ *x*_m,*i*_ − *X*_k_ (*i* = 1, …, *M*) and decouple the MT dynamics in a mean-field approximation, which neglects kinetochore velocity fluctuations.

We make two assumptions. First we assume that all *M* MTs are always attached to the kinetochore. Because the MT-kinetochore links are catch bonds this assumption is equivalent to assuming that these links are predominantly under tension. We will check below by comparison with numerical simulations to what extent this assumption can be justified. Secondly, we neglect that MTs are confined by a centrosome. Then, as mentioned above, the only relevant coordinates are the relative MT-kinetochore distances *x_i_*, which measure the extension of the *i*-th linker.

The MTs are coupled because they attach to the same kinetochore: each MT experiences a force *F*_mk,*i*_ = −*cx_i_* from the elastic linker to the kinetochore, which is under tension (compression) for *x_i_* < 0 (*x_i_ >* 0); the kinetochore is subject to the total counter force *F*_km_ = *c* ∑_*i*_ *x*_*i*_. Therefore, the kinetochore velocity *v*_k_ is a stochastic variable depending on *all* distances *x_i_*, on the one hand, but determines the velocities 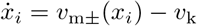 of MTs relative to the kinetochores, on the other hand. The equations can be decoupled to a good approximation because the one-sided system assumes a steady state with an approximately stationary kinetochore velocity *v*_k_ after a short time (rather than, for example, a cooperative oscillation as for an MT ensemble pushing against an elastic barrier [28]). In our mean-field approximation we then assume a constant kinetochore velocity *v*_k_ ≡ 〈*v*_k_〉 and neglect all stochastic fluctuations around this mean. This mean value is determined by the mean linker extension 〈*x*〉 via

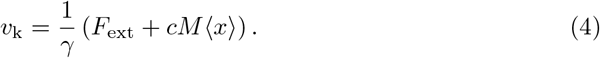

Fluctuations around the mean value are caused by fluctuations of the force *F*_km_ = *c* ∑_*i*_ *x*_*i*_ around its mean 〈*F*_km_〉 = *Mc* 〈*x*〉, which become small for large *M* (following the central limit theorem).

If *v*_k_ is no longer a stochastic variable, the dynamics of the MT-kinetochore extensions *x_i_* decouple. Then, the probability distribution for the *M* extensions *x_i_* factorizes into *M* identical factors *p*_±_(*x*_*i*_, *t*), where *p*_±_(*x, t*) are the probabilities to find one particular MT in the growing (+) or shrinking (−) state with a MT-kinetochore linker extensions *x*. We can derive two FPEs for the dynamical evolution of *p*_±_(*x, t*),

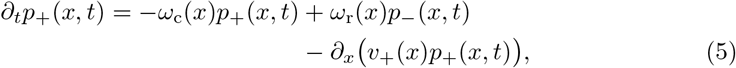

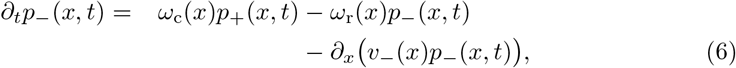

where *v*_±_(*x*) denotes the relative velocity of MT and kinetochore,

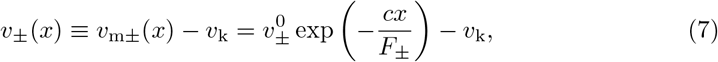

where *v*_k_ is no longer stochastic but self-consistently determined by Eq (4). We note that these FPEs are equivalent to single MT FPEs with position-dependent velocities, catastrophe and rescue rates [29–32].

We will now obtain the force-velocity relation of the whole MT ensemble by first solving the FPEs (7) in the stationary state *∂*_*t*_*p*_±_(*x, t*) = 0 and then calculating the mean linker extension 〈*x*〉 for given kinetochore velocity *v*_k_ using the stationary distribution *p*_±_(*x*). The external force that is necessary to move the kinetochore with velocity *v*_k_ then follows from Eq (4),

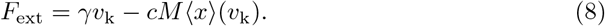

The MT-kinetochore distance *x* is limited to a maximal or a minimal value *x*_max_ or *x*_min_ for a given kinetochore velocity *v*_k_ > 0 or < 0, respectively, see Table 2. These limiting values are reached if the relative MT-kinetochore velocities vanish: If *v_k_* < 0 and MTs shrink, *x* shrinks until 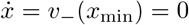 in Eq (7). If *v_k_* > 0 and MTs grow, *x* grows until 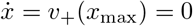. Linker extensions *x*_max_ (*x*_min_) are reached as stationary states if catastrophes (rescues) are suppressed (for example, because of large forces), such that MTs can grow (shrink) for sufficiently long times. If the external force *F*_ext_ is prescribed rather than a kinetochore velocity, all MTs reach a stationary state with the same velocity given by Eq (8). In this stationary state, both MT-tips and kinetochore move with the same velocity

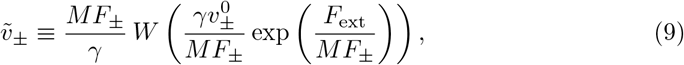

where *W* () denotes the Lambert-W function.

**Table 2.** Boundaries for linker extension. Maximal or a minimal value *x*_max_ or *x*_min_ of the stationary linker extension distribution *p*(*x*) from conditions *v*_−_(*x*_min_) = 0 and *v*_+_(*x*_max_) = 0.

|  |  |  |
| --- | --- | --- |
| $v_k > 0$ | $v_-(x) < -v_k$ always<br>$x_{\min} = -\infty$ | $v_+(x) > 0$ for $x < x_{\max}$<br>$x_{\max} = (F_+/c) \ln(v_+^0/v_k)$ |
| $v_k < 0$ | $v_-(x) < 0$ for $x > x_{\min}$<br>$x_{\min} = (F_-/c) \ln(v_-^0/v_k)$ | $v_+(x) > v_k$ always<br>$x_{\max} = \infty$ |
| $v_k = 0$ | $v_-(x) < 0$ always<br>$x_{\min} = -\infty$ | $v_+(x) > 0$ always<br>$x_{\max} = \infty$ |

In the complete absence of stochastic switching between growth and shrinking by catastrophes and rescues, the MT ensemble reaches stationary states with peaked distributions *p*_+_(*x*) ∝ *δ*(*x*_max_ − *x*) and *p*_−_(*x*) ∝ *δ*(*x* − *x*_min_). Stochastic switching shifts and broadens these peaks, and the FPEs (5) and (6) lead to a distribution *p*_±_(*x, t*) of linker extensions *x* in the growing and shrinking states with statistical weight *p*_±_(*x, t*) > 0 in the whole interval *x*_min_ ≤ *x* ≤ *x*_min_. At the boundaries *x*_min_ and *x*_max_ of this interval, the probability current density

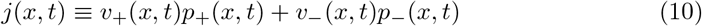

has to vanish. Furthermore, in any stationary state (*∂*_*t*_*p*_±_(*x, t*) = 0) the current density is homogeneous, as can be seen by summing up the FPEs (5) and (6):

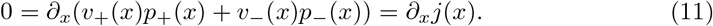

Together with *j* = 0 at the boundaries this implies that *j* = 0 everywhere in a steady state. The resulting relation *v*_+_(*x*)*p*_+_(*x*) = − *v*_−_(*x*)*p*_−_(*x*) can be used to reduce the stationary FPEs to a single ordinary differential equation with the solution [31]

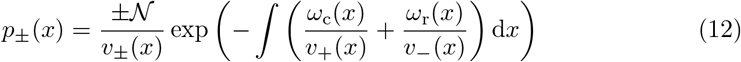

for the stationary distribution of linker extensions *x* in the growing and shrinking states. The normalization constant 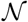 [must be chosen so that the overall probability density *p*(*x*) ≡ *p*_+_(*x*) + *p*_−_(*x*) satisfies 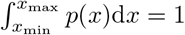. Obviously, *p*_±_(*x*) = 0 for *x* > *x*_max_ and *x* < *x*_min_. The stationary probability densities *p*_±_(*x*) from Eq (12) can then be used to calculate the mean distance 〈*x*〉 as a function of the kinetochore velocity *v_k_*, which enters through Eq (7) for *v*_±_(*x*). The integral in the exponent in Eq (12) as well as the normalization can be evaluated numerically to obtain an explicit 〈*x*〉 (*v*_k_)-relation, which is shown in Fig. 2A.

**Fig 2.**
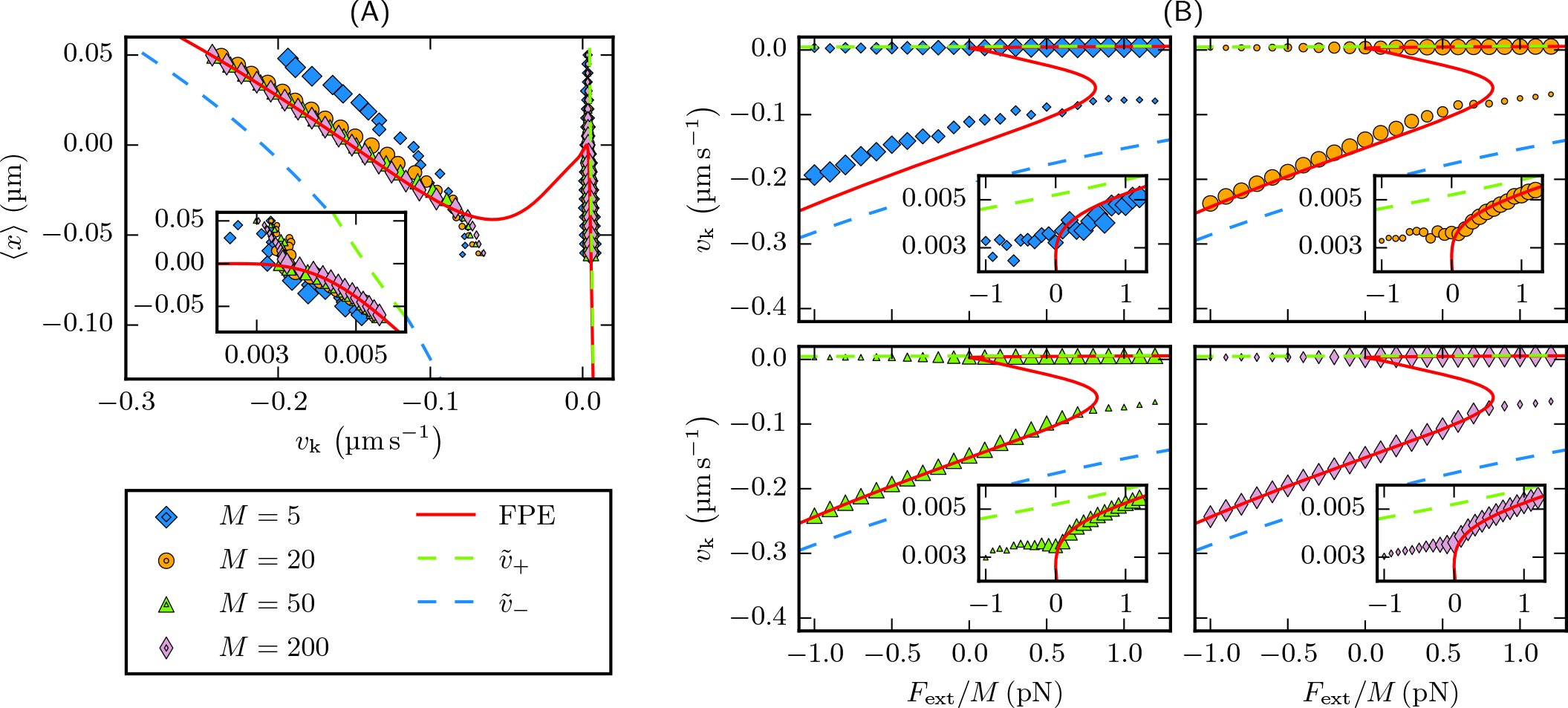
Mean-field results compared to stochastic simulations of the one-sided model. (A) The master curve 〈*x*〉(*v*_k_) from the mean-field approach (red line) agrees with simulation results for different MT-numbers *M* = 5, 20, 50, 200. The dashed lines mark *x*_min,max_(*v*_k_) from Table 2. We run simulations with constant external forces and average over 80 simulations for each force. Initially, the MT-kinetochore distance is either *x*_min_ or *x*_max_ while all MTs grow or shrink with velocity 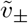, respectively. The system then enters a (meta-)stable state, in which we measure the mean kinetochore velocity and MT-kinetochore distances. The marker size depicts the time the system rests in this state on average, which is a measure for its stability (maximum marker size corresponds to *t*_rest_ ≥ 1000 s). As predicted, the mean-field approach turns out to be correct in the limit of many MTs, and in this limit the 〈*x*〉(*v*_k_)-relation is independent of the MT-number *M*. (B) Resulting force-velocity relations for different MT-numbers *M* = 5, 20, 50, 200. The dashed lines show the large velocity limit 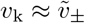 given by Eq (9). We used a linker stiffness of *c* = 20 pN μm^−1^ both in (A) and (B).

At this point it should be noticed that in the mean-field theory the *x* (*v*_k_)-relation is *independent* of the MT number *M*. Therefore, we call it *master curve* henceforth. In Fig. 2A we compare the mean-field theory result to stochastic simulations and find that the mean-field approach becomes exact in the limit of large *M*, where fluctuations in the kinetochore velocity around its mean in Eq (4) can be neglected.

The master curve is a central result and will be the basis for all further discussion. Together with the force-balance (8) on the kinetochore, the master curve will give the force-velocity relation for the MT-kinetochore system. A positive slope of the master curve, as it can occur for small negative *v*_k_ ≈ 0 (see Fig 2A), gives rise to an instability of the MT-kinetochore system: Then, a positive kinetochore velocity fluctuation *δv*_k_ > 0 leads to a MT-kinetochore linker compression *δ*〈*x*〉 > 0. According to the force-balance (8), a compression *δ*〈*x*〉 > 0 puts additional forward-force on the kinetochore leading to a positive feedback and further increase *δv*_k_ > 0 of the kinetochore velocity. This results in an instability, which will prevent the system to assume mean linker extensions 〈*x*〉 in this unstable regime. This is confirmed by stochastic simulation results in Fig 2A, which show that the unstable states are only assumed transiently for very short times. Therefore, occurrence of a positive slope in the master curve in Fig 2A is the essential feature that will give rise to bistability in the one-sided model and, finally, to oscillations in the full two-sided model.

Now we want to trace the origin of this instability for small negative *v*_k_ ≈ 0. If the MTs are growing (shrinking) for a long time, all linker extensions assume the stationary values *x* ≈ *x*_max_(*v*_k_) (*x* ≈ *x*_min_(*v*_k_)) from Table 2, where the MT-velocity adjusts to the kinetochore velocity, *v*_k_ ≈ *v*_m±_(*x*). If the kinetochore velocity increases in these states by a fluctuation (i.e., *δv*_k_ > 0), the MT-kinetochore linkers are stretched (i.e., *δx* < 0), which slows the kinetochore down again resulting in a stable response. This is reflected in the negative slopes of both *x*_max_(*v*_k_) (for *v*_k_ > 0) and *x*_min_(*v*_k_) (for *v*_k_ < 0). Because of constant stochastic switching between catastrophes and rescues the mean linker extension exhibits fluctuations about *x*_max_ and *x*_min_, but we expect also the master curve 〈*x*〉(*v*_k_) to have a negative slope for a wide range of velocities *v*_k_. Figure 2A shows that this is actually the case for kinetochore velocities *v*_k_ around the force-free growth or shrinking velocities 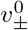 of the MTs, i.e., if the imposed kinetochore velocity *v*_k_ roughly “matches” the force-free growing or shrinking MT velocity. Then a small mismatch can be accommodated by small linker extensions *x*, which do not dramatically increase fluctuations by triggering catastrophe or rescue events.

The situation changes for small negative or small positive values of the kinetochore velocity around *v*_k_ ≈ 0. For *v*_k_ ≲ 0, MT-kinetochore linkers develop logarithmically growing large negative extensions *x*_min_ (see Table 2) corresponding to a slow kinetochore trailing fast shrinking MTs that strongly stretch the linker. Likewise, for *v*_k_ ≳ 0, MT-kinetochore linkers develop logarithmically growing large positive extensions *x*_max_ corresponding to a slow kinetochore trailing fast growing MTs that strongly compress the linker. Around *v*_k_ ≈ 0, the system has to switch from large negative *x* to large positive *x* because the resulting tensile force *F*_mk_ = −*cx* on the shrinking MT will destabilize the shrinking state and give rise to MT rescue at least for *x* < −*F_r_/c*.

Therefore, also the mean value 〈*x*〉 switches from negative to positive values resulting in a positive slope of the master curve if the stationary distributions *p*_−_(*x*) and *p*_+_(*x*) remain sufficiently peaked around the linker extensions *x*_min_ and *x*_max_, also in the presence of fluctuations by catastrophes and rescues. In S1 Appendix we show that the stationary distributions assume a power-law behavior *p*_+_(*x*) ∝ (*x*_max_ − *x*)^*α*+^ [*p*_−_(*x*) ∝ (*x* − *x*_min_)^*α*−^] around *x*_max_ [*x*_min_] for *v*_k_ > 0 [*v*_k_ < 0] with exponents *α*_±_ that depend on the MT-kinetochore stiffness *c* as *α*_±_ + 1 ∝ 1*/c* in the presence of fluctuations. It follows that distributions are peaked (i.e., have a large kurtosis) and bistability emerges if the MT-kinetochore linker stiffness *c* is sufficiently large such that deviations of the MT velocity from the kinetochore velocity become suppressed by strong spring forces. This is one of our main results. We also find that 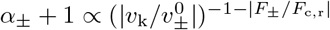 such that the distributions become also peaked around *x*_min,max_ in the of large velocities *|v*_k_*|*. Then the velocity approaches 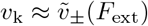 for a prescribed external force such that 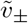 from Eq (9) represents the large velocity and large force limit of the force-velocity relation of the kinetochore (see Fig 2B).

In the unstable regime around *v*_k_ ≈ 0, the linker length distribution *p*(*x*) is typically broad without pronounced peaks and has a minimal kurtosis (as a function of *v*_k_) in the presence of catastrophe and rescue fluctuations. In this regime the system assumes a state with a heterogeneous stationary distribution of growing and shrinking MTs, i.e., the total probabilities to grow or shrink become comparable, ∫ *p*_+_(*x*)d*x* ∼ ∫ *p*_−_(*x*)d*x*. If the kinetochore velocity is increased, *δv*_k_ > 0, the system does not react by *δx* < 0, i.e., by increasing the average tension in the linkers in order to pull MTs forward, but by *switching* MTs from the shrinking to the growing state (on average), which then even allows to relax the average linker tension.

Using the force-balance (8) on the kinetochore, the master curve is converted to a force-velocity relation for the MT-kinetochore system; the results are shown in Fig 2B. The bistability in the master curve directly translates to a bistability in the force-velocity relation of the MT ensemble, and we obtain a regime with three branches of possible velocities for the same external force. The upper and the lower branches agree with our simulation results and previous simulation results in Ref. [24], and our mean-field results become exact in the limit of large *M*, see Fig 2B. These branches correspond to the two stable parts of the master curve with negative slope, that are found for kinetochore velocities *v*_k_ roughly matching the force-free growth or shrinking velocities 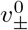 of the MTs. The mid branch corresponds to the part of the master curve with positive slope, where the system is unstable. Also Fig 2B demonstrates that this instability is confirmed by stochastic simulations results.

Finally, we note that a simpler theoretical approach, where it is assumed that all linkers assume *identical* extensions *x_i_* ≈ *x*, is exact for a single MT (*M* = 1) by definition but not sufficient to obtain a bistable force-velocity relation for MT ensembles (*M* > 1) (see S1 Appendix).

## 4 Oscillations in the two-sided model

As already worked out by Banigan *et al.* [24], the bistability in the force-velocity relation of the one-sided MT ensemble can be considered to be the cause for stochastic oscillations in the two-sided model. The external force in the one-sided model is a substitute for the spring force *F*_kk_ = *c*_k_ (*X*_k,r_ − *X*_k,l_ − *d*_0_) of the cohesin bond in the full model with a stiffness *c*_k_ and rest length *d*_0_, see Table 1. Since the cohesin force is a linear function of the inter-kinetochore distance, the force-velocity relation can be treated as phase space diagram for the two kinetochores (see Fig 3A), where both kinetochores move as points on the force-velocity relation. The cohesin bond always affects the two kinetochores in the same way because action equals reaction: if the cohesin spring is stretched, both kinetochores are pulled away from their pole (AP), if it is compressed, both kinetochores are pushed polewards (P). Thus, the kinetochores always have the same position on the *F*_kk_-axis in the *F*_kk_-*v*_k_ diagram in Fig 3A, if *F*_kk_ on the horizontal axis is defined as the force on the kinetochore in AP-direction (i.e., *F*_kk,l_ ≡ *F*_kk_ and *F*_kk,r_ ≡ −*F*_kk_ for the left/right kinetochore). Likewise, we define *v*_k_ on the vertical axis as the velocity in AP-direction (i.e., 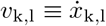 and 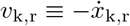 for the left/right kinetochore). The upper/lower stable branch of the force-velocity relation is denoted by 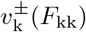. A kinetochore on the upper (lower) branch thus moves in AP-(P-)direction if 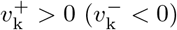, which is the typical situation. Using *F*_kk_ = −*c*_k_ (*X*_k,r_ − *X*_k,l_ − *d*_0_) for the spring force, we find 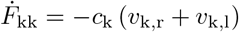, i.e., kinetochores move with the sum of their AP-velocities along the force-velocity curve in the *F*_kk_-*v*_k_ diagram.

**Fig 3.**
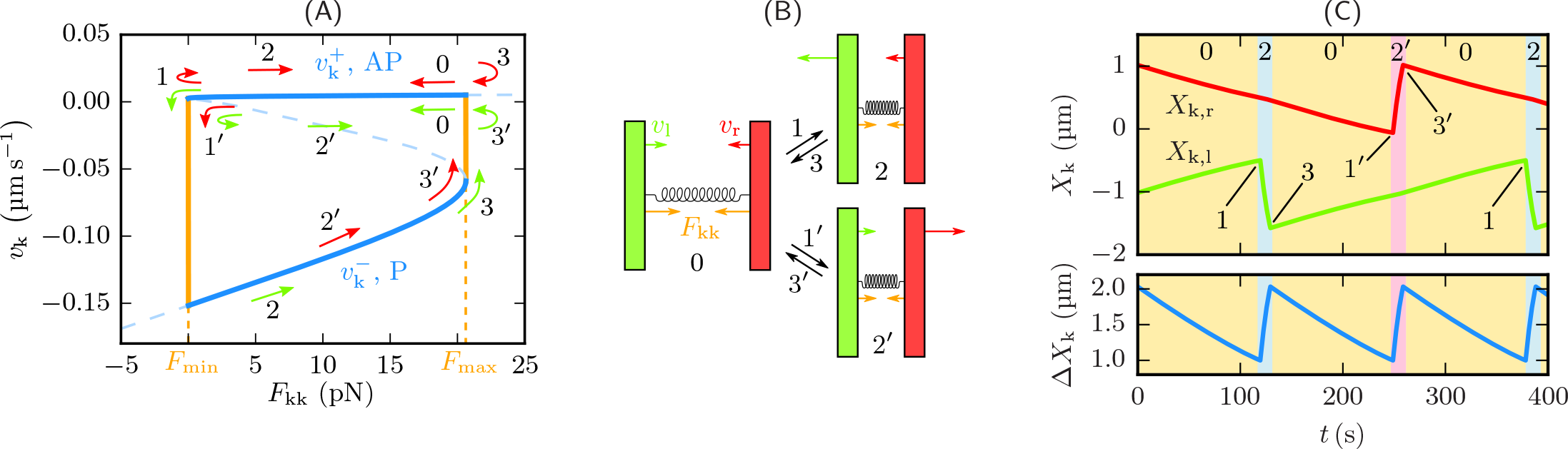
Bistability gives rise to oscillations in the two-sided model. (A,B) Different states of sister kinetochore motion can be deduced from the bistability of the force-velocity relation: either both kinetochores are in the upper branch (0) or one is in the upper and the other one in the lower branch (2, 2′). In the first case, both kinetochores move away from their pole (AP) towards each other. Thus, the spring force *F*_kk_ decreases until it reaches *F*_min_. Since the upper branch is not stable anymore below *F*_min_, either the left (1) or the right (1′) kinetochore switches to the lower branch and changes direction to poleward movement (P). The system is then in state 2 or 2′, where both kinetochores move into the same direction: the leading kinetochore P, the trailing kinetochore AP. As P-is much faster than AP-movement (due to difference between MT shrinking and growth velocities), the inter-kinetochore distance and the spring force are increasing. Above *F*_max_ only AP-movement is stable, which is why the leading kinetochore changes direction (3, 3′) and the system switches to state 0 again. (C) Solution of the equations of motion (13) for *c* = 20 pN μm^−1^ and *M* = 25. The initial condition is *F*_kk_ = *F*_max_ (both kinetochores at the right end of the upper branch). For an animated version see S1 Video.

Oscillations arise from the two kinetochores moving through the hysteresis loop of the bistable force-velocity relation as described in Fig 3A. Three states are possible (see Fig 3B). In state 0, both kinetochores move in AP-direction (i.e., in opposite directions) relaxing the *F*_kk_-force from the cohesin bond, i.e., on the upper branch and to the left in the *v*_k_-*F*_kk_-diagram with velocity 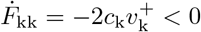. After reaching the lower critical force *F*_min_ of the hysteresis loop, one of the two kinetochores reverses its direction and switches to the lower branch resulting into states 2 or 2′ where one kinetochore continues in AP-direction with 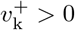 while the other is moving in P-direction with 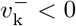 (i.e., both move in the same direction). In the *v*_k_-*F*_kk_-diagram, this results in a motion to the right with velocity 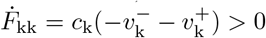 because MTs typically shrink much faster than they grow (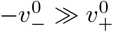, see Table 1). Moving on opposite P- and AP-branches increases the kinetochore distance and builds up *F*_kk_-force in the cohesin bond. After reaching the upper critical force *F*_max_ of the hysteresis loop, it is always the kinetochore on the lower branch moving in P-direction which switches back and state 0 is reached again. This behavior is in agreement with experimental results [11]. The system oscillates by alternating between state 0 and one of the states 2 or 2′ (which is selected randomly with equal probability).

For each of the states 0, 2 and 2′depicted in Fig 3AB the two branches 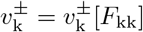 provide deterministic equations of motion for the kinetochore positions. Inserting *F*_kk_ = −*c*_k_ (*X*_k,r_ − *X*_k,l_ − *d*_0_) we obtain both kinetochore velocities as functions of the kinetochore positions and find

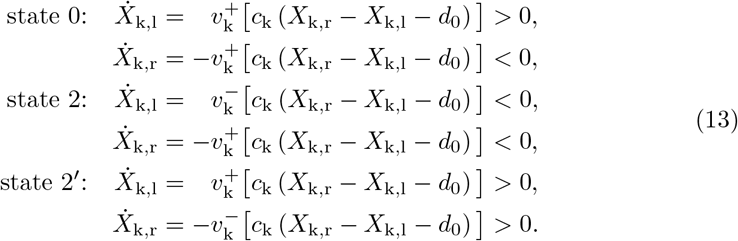

Solving these equations gives idealized deterministic trajectories of the sister kinetochores, when we also assume that the left and the right kinetochore pass the lower branch alternately such that the order of states is a periodic sequence 0 − 2 − 0 − 2′ −0 … as shown in the example in Fig 3C. Then single kinetochores oscillate with half the frequency of inter-kinetochore (breathing) oscillations, just as observed in PtK1 cells [11]. Moreover, we can obtain numerical values of the frequencies directly from the trajectories. For a MT-kinetochore linker stiffness *c* = 20 pN μm^−1^ and 20–25 MTs per kinetochore, which is a realistic number for mammalian cells [33], we get periods of 206–258 s and 103–129 s for kinetochore and breathing oscillations, respectively. These values coincide with experimental results of 239 s and 121 s measured in PtK1 cells [11].

The calculated trajectories are idealized since they neglect stochastic fluctuations that occur in simulations of the two-sided model and have two main effects on the kinetochore dynamics. Firstly, the sister kinetochores do not pass the lower branch alternately but in random order. Therefore, we observe phases where one kinetochore moves in AP-direction for several periods, while the other one changes its direction periodically but moves polewards on average (Fig 4A). Since this does not influence the trajectory of the inter-kinetochore distance, breathing oscillations still occur in a more or less regular manner, which allows us to measure their frequencies by Fourier analysis. We will show below that additional polar ejection forces suppress this random behavior and force the kinetochores to pass the lower branch alternately. As a second effect of the stochastic character of the simulation, kinetochores do not change the branch instantaneously after crossing the critical forces *F*_max_ or *F*_min_. Instead, they tend to maintain their primary state for a while (Fig 4B) and follow the metastable states that we also observe in the one-sided model (Fig 2B). Hence, the frequencies we measure in the simulations are smaller than those we calculate from the Fokker-Planck mean-field approach (Fig 4C). The latter effect vanishes in the limit of many MTs (large *M*): the switching points approach the theoretical values *F*_max_ and *F*_min_, and the simulated breathing frequencies converge to our mean-field predictions.

**Fig 4.**
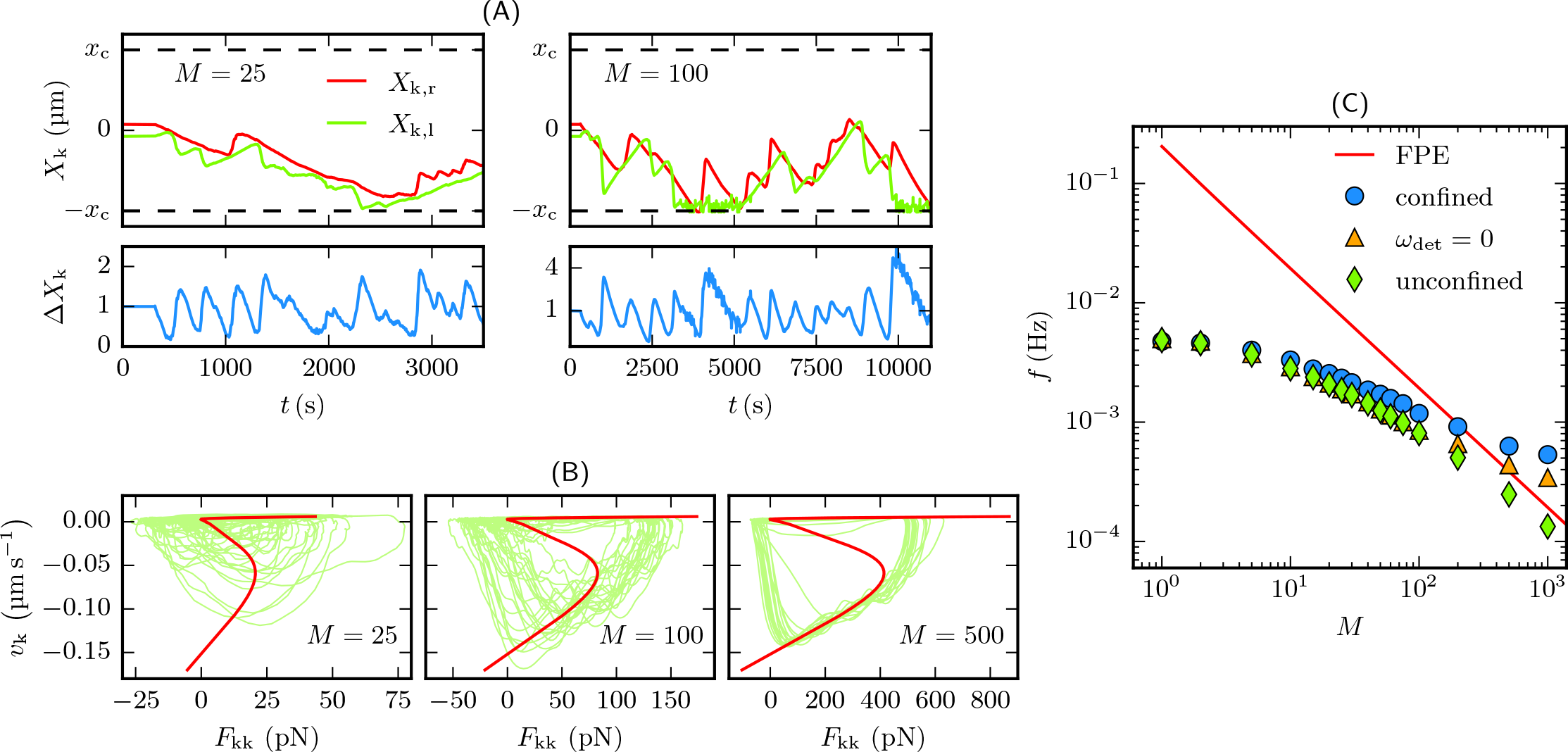
Oscillations in stochastic simulations compared to mean-field results. (A) Kinetochore trajectories and breathing oscillations in the confined two-sided model as depicted in Fig 1A. There are phases where one kinetochore moves more or less constantly AP while the other one oscillates. The breathing oscillations are regular enough to assign a frequency by Fourier analysis. With 100 MTs one kinetochore can get stuck to the centrosome for a while. (B) Kinetochore velocity against cohesin force in simulations of the two-sided model without confinement (*x*_c_ → ∞) and detachment (*ω*_d_ = 0) (green). For many MTs the velocity follows very precisely the predicted hysteresis from the mean-field approach (red). For animated versions see S2 Video (*M* = 25) and S3 Video (*M* = 500). (C) Double-logarithmic plot of frequencies of breathing oscillations as a function of MT number *M*: calculated from the mean-field approach according to Fig 3 (red) and measured in simulations of the unconfined (green diamonds) as well as the confined model with (blue circles) and without detachment (orange triangles). Confinement becomes relevant for large MT numbers. In the presence of detachment only 75 % of the MTs are attached on average, which corresponds to a simple shift of the curve to lower MT numbers. For all simulations the MT-kinetochore linker stiffness was *c* = 20 pN μm^−1^.

In the limit of large *M*, on the other hand, confinement by the centrosome influences the kinetochore dynamics: since more MTs exert a higher force on the kinetochore, it is possible that one of the two sisters gets stuck at the centrosome for a while (see Fig 4A). Hence, the frequencies measured in the confined two-sided model deviate from the frequencies in the unconfined case above *M* ≈ 200.

We suppress detachment in simulations of the unconfined model to prevent unattached MTs from shrinking towards infinity. In the confined model, dynamics does qualitatively not depend on whether the MTs are able to detach from the kinetochore, i.e., to rupture the catch bond. In particular, the occurrence of oscillations is robust against inclusion of MT-kinetochore detachment. If we enable detachment in our simulations we find that, on average, about 75 % of the MTs are attached independently of the total MT number. Therefore, such simulations behave effectively as a system without detachment but with less MTs, which explains the difference in frequencies between the confined models with and without detachment in Fig 4C.

## 5 Constraints for bistability and oscillations

### 5.1 Constraints for bistability in the one-sided model

We already argued above in Sec. 3 that bistability (and thus oscillations) can only emerge if the MT-kinetochore linker is sufficiently stiff. To analyze the influence of the linker stiffness *c* and the MT number *M* on bistability quantitatively, the transformation from the master curve to the force-velocity relation is visualized in Fig 5A as search for the intersections of the master curve with linear functions

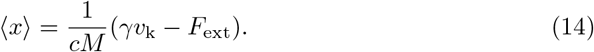

In the limit of large *M* these linear functions have zero slope. Bistable force-velocity relations with three intersection points are only possible if the master curve has positive slope for intermediate *v*_k_ resulting in a maximum and minimum. The extrema of the master curve vanish, however, in a saddle-node bifurcation if the linker stiffness drops below *c*_bist_ = 7.737 pN μm^−1^, which is, therefore, a lower bound for the occurrence of bistability. In the case of finite MT numbers *M*, bistable force-velocity relations can only be found if the slope in the inflection point of the master curve exceeds *γ/cM* (the slope of the linear function (14)). This allows us to quantify a bistable regime in the parameter plane of linker stiffness *c* and MT number *M* as shown in Fig 5B.

**Fig 5.**
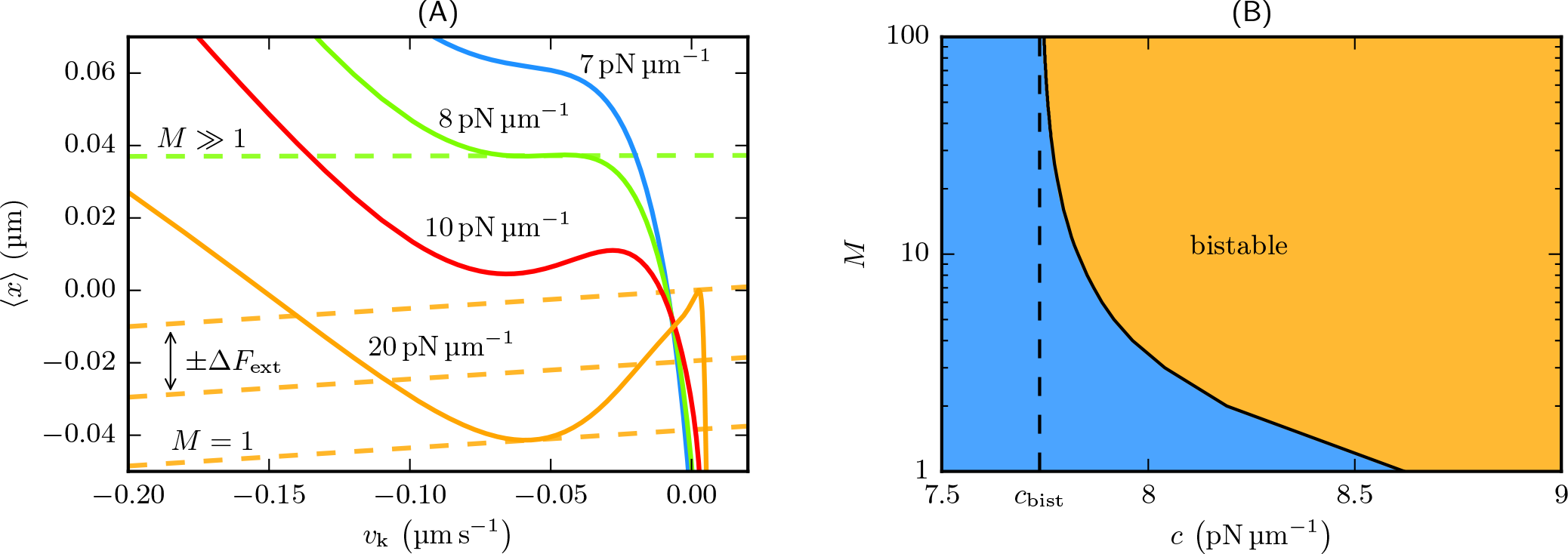
Constraints for bistability in the one-sided model. (A) Master curves for different linker stiffnesses *c* and linear functions according to (14). In the limit of large *M* the linear functions have zero slope and bistability occurs if the master curve has two extrema, which is the case for *c* > *c*_bist_. For finite *M* bistable solutions are possible if the linear functions have a smaller slope than the inflection point of the master curve. (B) Resulting bistable regime in the parameter plane of linker stiffness *c* and MT number *M*.

### 5.2 Constraints for oscillations in the two-sided model

We showed in Sec. 4 that bistability of the one-sided model is a necessary condition for oscillations in the two-sided model. Now we show that bistability in the one-sided model is, however, *not sufficient* for oscillations in the full model. If the force-velocity relation is interpreted as phase space diagram for the two kinetochores, kinetochores only switch branches in the *v*_k_-*F*_kk_-diagram if their velocity changes its sign at the turning points *F*_min_ and *F*_max_. If this is not the case and one of the two branches crosses *v*_k_ = 0 (e.g. the right branch for *c* = 10 pN μm^−1^ in Fig 5A, which transforms to the upper branch of the force-velocity relation), the intersection point is a stable fixed point in the phase space diagram (see Fig 6A). At this fixed point kinetochore motion will relax to zero velocity and just exhibit fluctuations around an equilibrium distance instead of oscillations.

**Fig 6.**
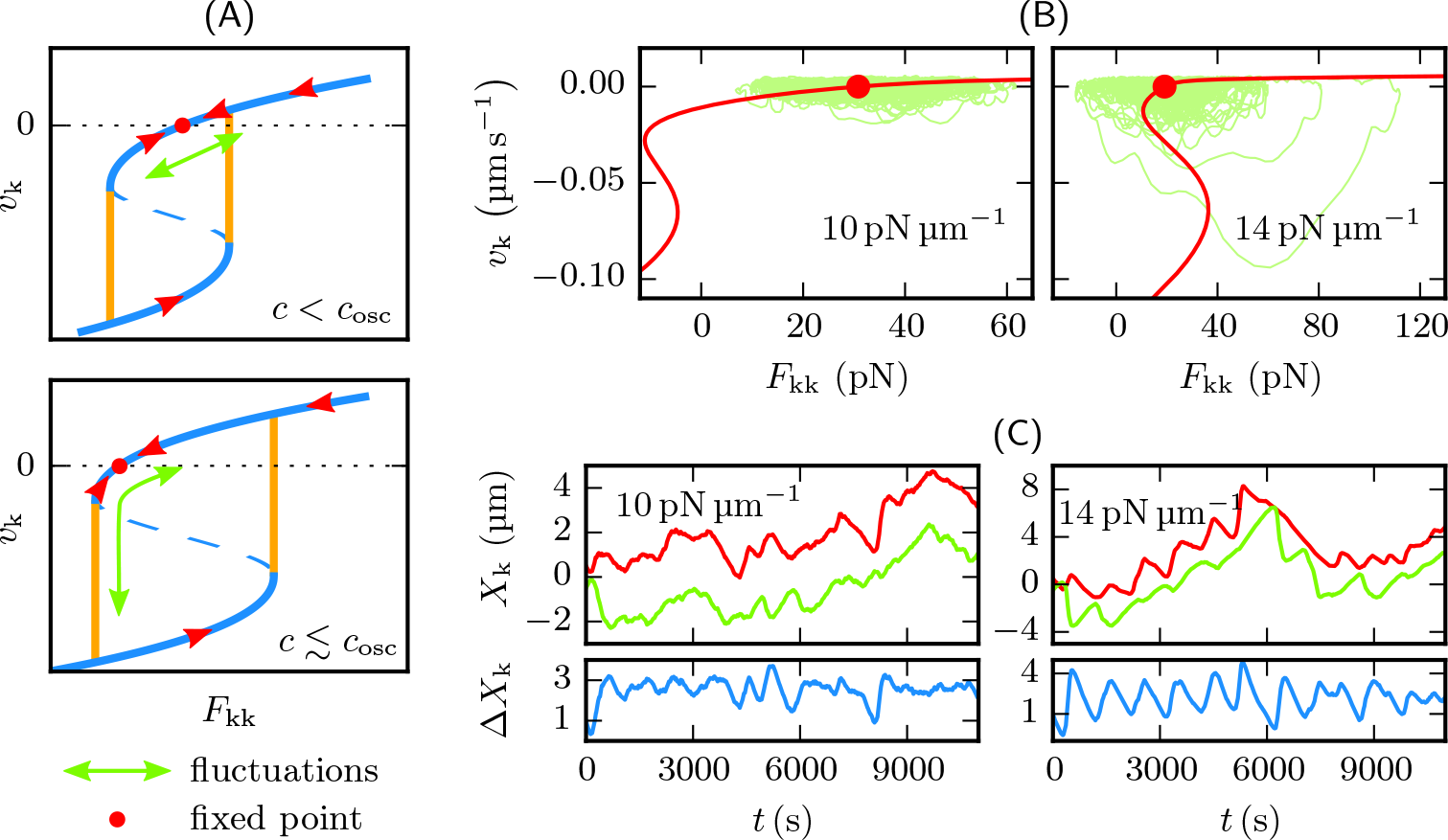
Kinetochore dynamics in the non-oscillatory regime. (A) Schematic explanation of kinetochore motion in the non-oscillatory regime based on the force-velocity relation. Where the upper branch crosses zero velocity, inter-kinetochore distance has a fixed point, around which it fluctuates. With higher linker stiffnesses *c* the fixed point comes closer to the left turning point *F*_min_. When *c* is just slightly smaller than *c*_osc_, fluctuations can be large enough for the kinetochore distance to leave the upper stable branch. Then, one of the two sister kinetochores passes once through the lower branch. (B,C) This behavior can be observed in simulations. While at *c* = 10 pN μm^−1^ kinetochores just fluctuate around the fixed point, at *c* = 14 pN μm^−1^ the kinetochores occasionally pass through the hysteresis loop. Simulations were performed with an unconfined system and 100 MTs on each side.

As a sufficient condition for oscillations we have to require – besides bistability – a strictly positive velocity in the upper and a strictly negative velocity in the lower branch in the *v*_k_-*F*_kk_-diagram. Based on this condition we quantify an oscillatory regime in the parameter plane of linker stiffness *c* and MT number *M* in Fig 7A. In the limit of many MTs the sufficient condition for oscillations can be formulated in terms of the master curve: the maximum of the master curve has to be located at a positive and the minimum at a negative velocity. This is the case for *c* > *c*_osc_ = 15.91 pN μm^−1^, which is, therefore, a lower bound for the occurrence of oscillations. This constraint on the linker stiffness for metaphase chromosome oscillations provides additional information on MT-kinetochore linkers whose molecular nature is not known up to now.

**Fig 7.**
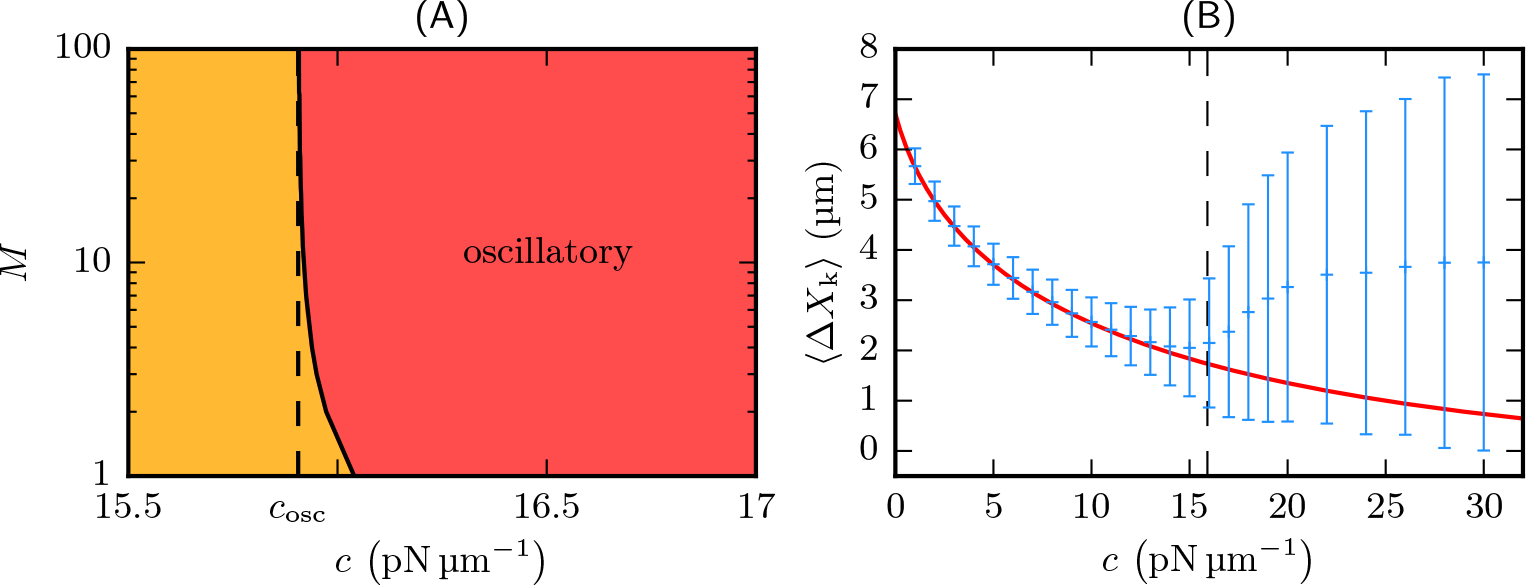
Constraints for oscillations in the two-sided model. (A) Oscillatory regime in the parameter plane of linker stiffness *c* and MT number *M*. (B) Mean inter-kinetochore distance according to Eq (16) (red) and measured in simulations (blue) with *M* = 100. Below *c*_osc_ = 15.91 pN μm^−1^ (dashed line) both results match, whereas in the oscillatory regime mean inter-kinetochore distance diverges from the fixed point, and its standard deviation increases notably.

Because of stochastic fluctuations, the transition between oscillatory and non-oscillatory regime is not sharp in our simulations. In the non-oscillatory regime kinetochores fluctuate around a fixed point of inter-kinetochore distance, where the upper branch crosses *v*_k_ = 0. However, these fluctuations can be large enough for the inter-kinetochore distance to shrink and leave the upper branch on the left side, especially for stiffnesses *c* slightly below *c*_osc_. If that happens, one kinetochore passes once through the lower branch of the force-velocity relation just as in an oscillation. The difference to genuine oscillations is that these are randomly occurring single events (resulting in a Poisson process). Randomly occurring oscillations are visualized in Fig 6 for *c* < *c*_osc_ and *c* ≲ *c*_osc_. Moreover, the force-velocity relations as well as the kinetochore trajectories measured in corresponding simulations are shown.

In the non-oscillatory regime, the fixed point should determine the mean inter-kinetochore distance 〈∆*X*_k_〉 = 〈*X*_k,r_ − *X*_k,l_〉. Solving the FPEs for *v*_k_ = 0, we compute the (external) force *F*_0_ that has to be applied to one kinetochore to stall its motion:

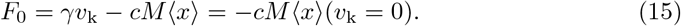

In the two-sided model this force is applied to the kinetochores by the cohesin bond at the fixed point. With *F*_kk_ = *c*_k_(∆*X*_k_ − *d*_0_) we compute the corresponding mean inter-kinetochore distance:

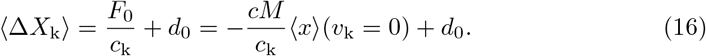

Figure 7B shows that simulations agree with this result in the non-oscillatory regime. At *c*_osc_ the transition to the oscillatory regime can be recognized, where the mean inter-kinetochore distance deviates from the fixed point (16). Moreover, the variance of ∆*X*_k_ increases significantly at *c*_osc_ due to the transition to the oscillatory regime.

## 6 Poleward microtubule flux

An effect we have not included so far is poleward microtubule flux, which was observed in several metazoan cells (Table 3). It describes the constant flux of tubulin from the plus-ends towards the spindle pole and is probably driven by plus-end directed kinesin-5 motors pushing overlapping antiparallel MTs apart as well as kinesin-13 proteins that are located at the centrosome and depolymerize the MTs at their minus-ends [27]. During metaphase, spindle and MT length can be maintained by simultaneous polymerization at the plus-ends [34], which results in a behavior similar to treadmilling of MTs [35].

**Table 3.** Metaphase poleward flux velocities *v*_f_ and occurrence of directional instability. For a more detailed review of poleward flux measurements see Ref. [34]

| Cell type | $v_f/\text{nm s}^{-1}$ | Directional instability |
| --- | --- | --- |
| LLC-PK1 (porcine) | 8.3 [8] | yes [8] |
| PtK1 (rat kangaroo) | 7.7 [36] | yes [11] |
| PtK2 (rat kangaroo) | 10 [8] | yes [12] |
| Newt lung | 9.0 [37] | yes [6] |
| U2OS (human) | 8.8 [9] | yes [9] |
| Drosophila embryo | 32 [38] | no [13] |
| Xenopus egg | 37 [39] | no [14] |

Poleward flux can be easily included in our model by subtracting a constant flux velocity *v*_f_ from the MT velocity. Then, the relative MT-kinetochore velocities (7) become

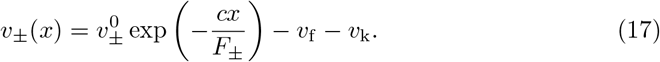

Hence, the flux velocity can be treated as an offset to the constant kinetochore velocity in the solution of the stationary FPEs. The final effect is a shift of both the master curves and the force-velocity relations by *v*_f_ towards smaller kinetochore velocities *v*_k_. If the shift is so large that the left turning point *F*_min_ of the force-velocity hysteresis is located at a negative velocity, poleward flux suppresses directional instability because a fixed point emerges, and we expect similar behavior as for intermediate linker stiffnesses in the previous section (see Fig 6). In the limit of many MTs, the maximum flux velocity that still allows directional instability is given by the velocity in the maximum of the master curve, which provides the boundary of the oscillatory regime in the parameter plane of linker stiffness *c* and poleward flux velocity *v*_f_ (Fig 8A). Phase space diagrams (Fig 8B) and kinetochore trajectories (Fig 8C) from simulations with appropriate flux velocities confirm our arguments exhibiting similar behavior as for intermediate linker stiffnesses in Fig 6. For small flux velocities the boundary of the oscillatory regime in Fig 8A approaches our above result *c*_osc_ = 15.91 pN μm^−1^. For increasing flux velocities the oscillatory regime shrinks, and its boundary has a maximum at *c* ≈ 50 pN μm^−1^ with *v*_f_ ≈ 3.11 nm s^−1^. We conclude that kinetochore oscillations can be suppressed by moderate flux velocities independently of the linker stiffness.

**Fig 8.**
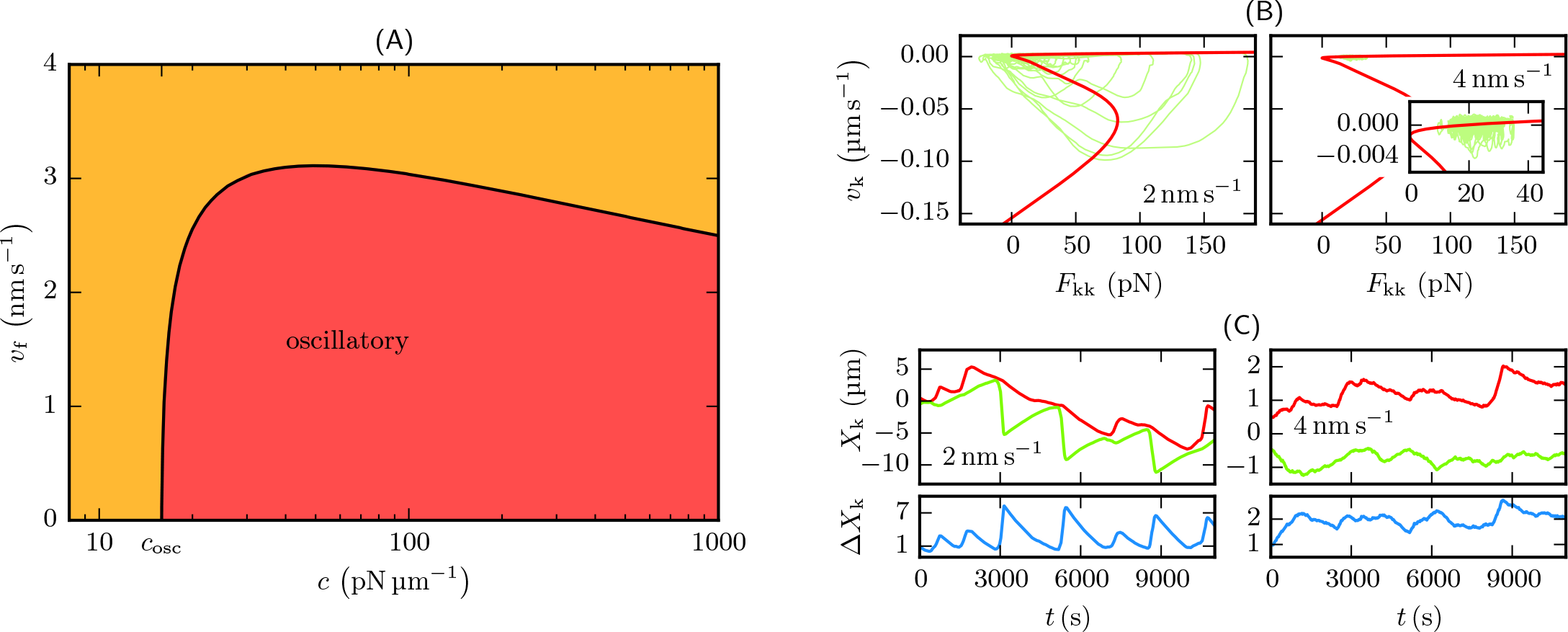
Poleward flux suppresses oscillations. (A) Oscillatory regime in the parameter plane of *c* and *v*_f_ in the limit of many MTs. Fast poleward flux suppresses kinetochore oscillations. (B,C) Phase space diagrams and MT trajectories from simulations of the unconfined two-sided model with *c* = 20 pN μm^−1^ and *M* = 100. While at *v*_f_ = 2 nm s^−1^ the system is still in the oscillatory regime, where hysteresis is recognizable in phase space, at *v*_f_ = 4 nm s^−1^ kinetochores show fluctuative motion as described in Fig 6.

Our theory also agrees with and explains simulation results in Ref. [24], where, for large flux velocities, suppression of kinetochore oscillations were observed but at the same time maintenance of bistability. Moreover, our results explain the experimentally observed correlation between flux velocity and directional instability. Kinetochore oscillations have been observed in the mitotic vertebrate cells listed in Table 3 (LLC-PK1, PtK1/2, newt lung, U2OS) which have poleward flux velocities not exceeding 10 nm s^−1^, whereas in the mitosis of a Drosophila embryo as well as in meiosis of a Xenopus egg, where flux velocities are three to four times higher, chromosomes do not exhibit directional instability.

## 7 Polar ejection forces

So far, we have not included polar ejection forces (PEFs). They originate from non-kinetochore MTs interacting with the chromosome arms and pushing them thereby towards the spindle equator, either through collisions with the chromosome arms or via chromokinesins [26], and provide additional pushing forces on kinetochores. Therefore, they can be included into the model by adding forces *F*_PEF,r_(*X*_k,r_) and *F*_PEF,l_(*X*_k,l_) acting on kinetochores, which depend on the *absolute* position of the kinetochores [23]. Due to the exponential length distribution of free MTs as well as the spherical geometry of the MT asters, the density of non-kinetochore MTs decreases monotonically with the distance from the spindle pole. Therefore, we assume that PEFs reach their maximum at the centrosome and vanish at the spindle equator (located at *x* = 0), where opposite PEFs compensate each other. We will only discuss linearized PEFs

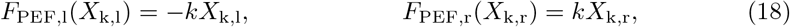

where the spring constant *k* defines the strength of the forces, and the signs are chosen so that a positive force acts in AP-direction. We checked that other more realistic force distributions do not differ qualitatively in their influence to the kinetochore dynamics.

To determine kinetochore trajectories of the two-sided model in the presence of PEFs, we can start from the same force-velocity relations as for the basic one-sided model. In the presence of PEFs, the total forces *F*_k,l_ and *F*_k,r_ that act on the left and the right kinetochore in AP-direction depend on the absolute kinetochore positions *X*_k,l_ and *X*_k,r_:

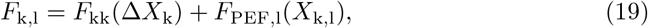

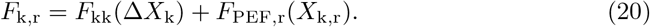

We can investigate the motion of kinetochores in the full two-sided model again by using a phase space diagram; in the presence of PEFs we use a *v*_k_-*F*_k_-diagram with the total force *F*_k_ in AP-direction on the horizontal axis and the velocity *v*_k_ in AP-direction on the vertical axis. Because the total forces contain the external PEFs they are no longer related by action and reaction and, thus, the two kinetochores no longer have the same position on the *F*_k_-axis, but they still remain close to each other on the *F*_k_-axis as long as the cohesin bond is strong enough.

A kinetochore on the upper/lower branch moves in AP-/P-direction with 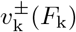 if 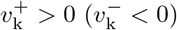. A kinetochore on the upper AP-directed branch will relax its AP-directed PEFs, while a kinetochore on the lower P-directed branch will build up AP-directed PEFs. After a time of equilibration the kinetochores behave as described in Fig 9. When one kinetochore changes its direction from P to AP (switches to the upper branch) the sister kinetochore, which was on the upper branch before, becomes the leading kinetochore (here, “leading” refers to the position in the force velocity phase space). Therefore, the kinetochores do not reach the left turning point *F*_min_ at the same time so that it is always the leading kinetochore that switches to the lower branch. Since in general the absolute P-velocity is much larger than the AP-velocity (−*v*_−_ for the lower branch is much larger than +*v*_+_ for the upper branch), the AP-directed PEF contribution to the total force increases faster on the lower branch than on the upper one. As a result, the P-moving kinetochore overtakes its sister on the *F*_k_-axis before switching back to the upper branch such that the leading kinetochore automatically becomes the trailing kinetochore in the next oscillation period (again, “leading” and “trailing” in terms of phase space positions). This periodic change of kinetochore positions in the force-velocity diagram leads to both regular breathing and regular single kinetochore oscillations, as the kinetochores alternately pass the lower branch. Solving appropriate equations of motions similar to Eqs (13) for each of the states depicted in Fig 9AB, we determine the deterministic trajectories in Fig 9C confirming this regular alternating oscillation pattern.

**Fig 9.**
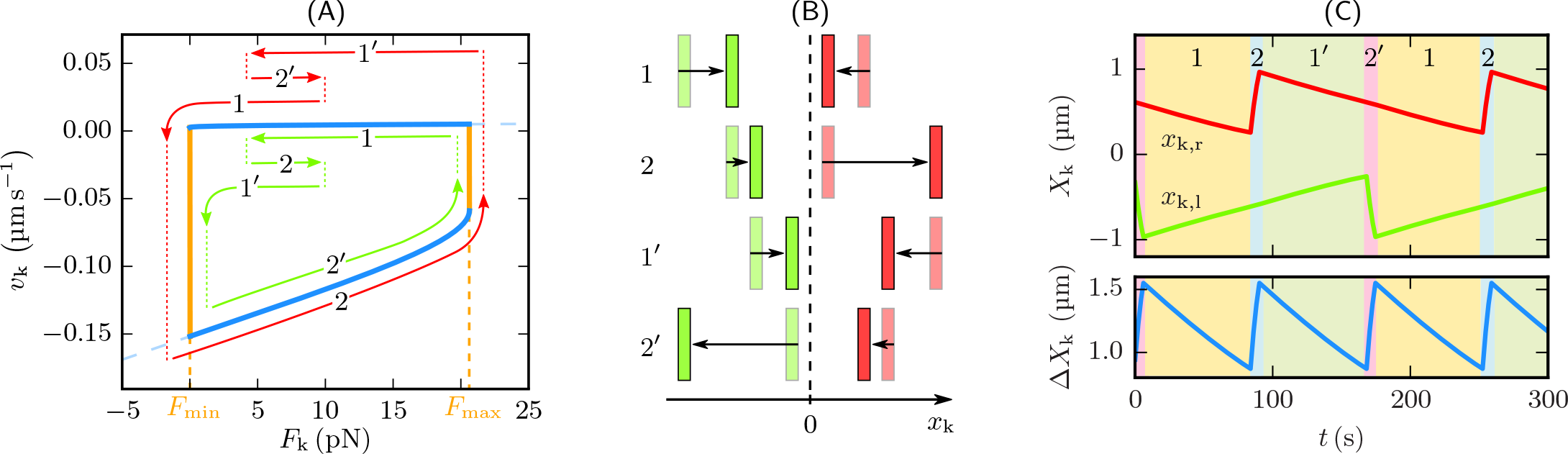
Kinetochore motion in the presence of PEFs. (A,B) At the beginning of state 1 the left kinetochore (green) has just switched from P- to AP-movement, so that both kinetochores are on the upper branch. Both kinetochores move in AP-direction, which means that both the cohesin force and the PEFs decrease and both kinetochores move left in the force-velocity diagram. Due to different PEFs, the right kinetochore (red) reaches the left turning point *F*_min_ first and switches to the lower branch, which marks the start of state 2. This state is dominated by the fast P-movement of the right kinetochore, which causes a steep increase of both *F*_kk_ and *F*_PEF,r_. Therefore, the right kinetochore moves to the right in the force-velocity diagram. Meanwhile, the left sister still moves in AP-direction and *F*_k,l_ increases slightly as the increase of *F*_kk_ is larger than the decrease of *F*_PEF,l_. Since 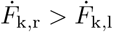, the right kinetochore overtakes its sister on the *F*_k_-axis before it reaches the right turning point and switches to the upper branch. The then following states 1′ and 2′ are the exact opposite to 1 and 2 with swapped kinetochores. (C) Solution of the corresponding equations of motion for *c* = 20 pN μm^−1^, *k* = 10 pN μm^−1^ and *M* = 25. For an animated version see S4 Video.

The alternating oscillation pattern in the presence of PEFs robustly survives in stochastic simulations as we demonstrate in Fig 10. Consequently, the kinetochores oscillate around the spindle equator and can not get stuck to one of the centrosomes as in the basic model, which means that PEFs are necessary to assure proper chromosome alignment in the metaphase plate at the spindle equator. This is consistent with an experiment by Levesque and Compton [40], who observed mitosis of vertebrate cells after suppressing the activity of chromokinesins and, thus PEFs. This resulted in 17.5 % of the cells in at least one chromosome not aligning at the equator, but locating near a spindle pole.

**Fig 10.**
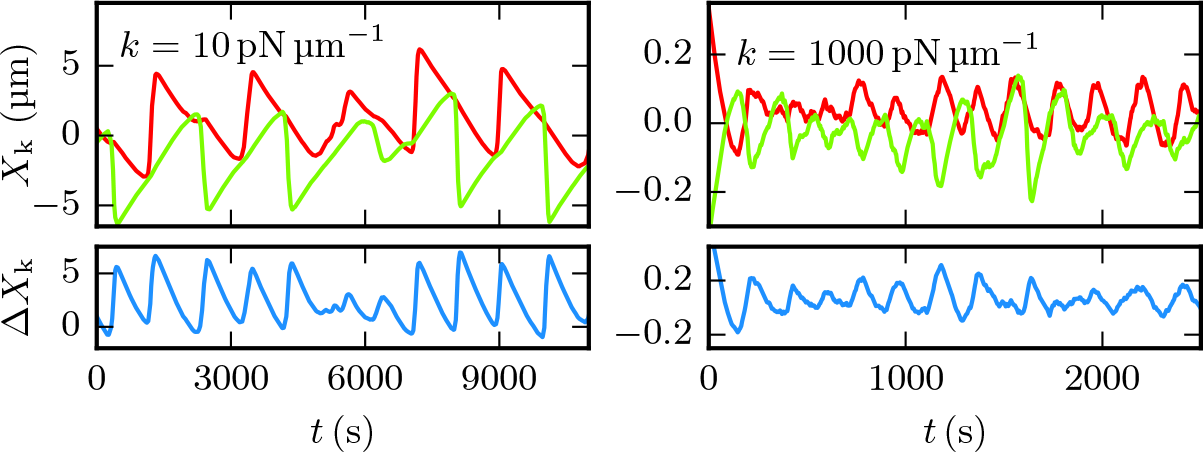
PEFs in stochastic simulations. Kinetochore trajectories with different PEF constants *k* from simulations with *M* = 100, *c* = 20 pN μm^−1^ and without confinement at the spindle poles. The PEFs force the kinetochores to oscillate regularly and to stay near the spindle equator. Contrary to the results of the FPE approach, stronger PEFs cause a more fluctuative kinetochore motion. Especially in systems with moderate MT numbers, this can lead to suppression of kinetochore oscillations. For animated versions of phase space trajectories see S5 Video (*k* = 10 pN μm^−1^) and S6 Video (*k* = 1000 pN μm^−1^).

Moreover, PEFs reduce the amplitude and increase the frequency of oscillations. The amplitude decreases for increasing PEF strength *k* as the kinetochores have to cover a smaller distance between the turning points at *F*_min_ and *F*_max_. The increase of the frequency is linear in *k*, which can be deduced from the linear increase of 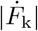:

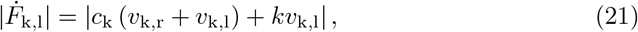

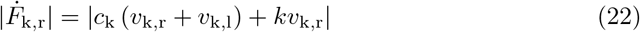

(defining 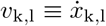 and 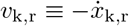 as the velocities in AP-direction as before). Since PEFs do not have any influence on the underlying master curves and force-velocity relations, they never completely suppress kinetochore oscillations in the deterministic Fokker-Planck model, but only reduce their amplitude and increase their frequency. For strong PEFs, however, this gives rise to kinetochore motion with a fluctuative character, see Fig 10 (see also S6 Video). The same observation was made in the model of Civelekoglu-Scholey *et al.* [23].

## 8 Microtubule confinement at the kinetochore

So far, we assumed that MTs are also able to exert pushing forces on the kinetochore. During oscillations we find, on average, slightly less (48%) MT-kinetochore links under tension, while a substantial part of linkers also exerts pushing forces. Two experimental results suggest, however, that MTs do not exert pushing forces: In Ref. [7], it was shown that the link between chromosomes is always under tension; the experiments in Ref. [25] demonstrated that, after removal of the cohesin bond, AP-moving kinetochores immediately stop because MTs can not exert pushing forces, while P-moving kinetochores continue moving due to MT pulling forces.

In view of these experimental results and in order to answer the question whether MT pushing forces are essential for bistability and oscillations, we analyze variants of our basic model, where MT growth is confined at the kinetochore, i.e., where the relative coordinate *x = x*_m_ − *X*_k_ is limited to *x* ≤ 0 such that MTs can only exert tensile forces on the kinetochore. This implies that the kinetochore undergoes a catastrophe if it reaches the kinetochore, i.e., if the relative coordinate reaches *x* = 0 from below in the one-sided model. Different choices for the corresponding catastrophe rate 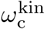 at *x* = 0 are possible: (i) A reflecting boundary, i.e., 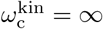, where a catastrophe is immediately triggered if the MT plus-end reaches the kinetochore. (ii) A “waiting” boundary condition, where the *relative* velocity *v*_+_ = *v*_m+_ − *v*_k_ = 0 stalls if the MT reaches *x* = 0 (in the simulation, we set the MT velocity to *v*_m+_ = *v*_k_). In contrast to the reflecting boundary condition, the catastrophe rate 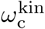 at the kinetochore is finite such that the MT waits at the kinetochore until it undergoes a catastrophe for a mean waiting time 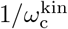. Because *x* = 0 also results in *F*_mk_ = 0, the force-free catastrophe rate seems a natural choice, 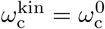 [see Eq (1)], which should be realized in the absence of any additional catastrophe regulating proteins at the centromere. (iii) If catastrophes are promoted by regulating proteins, but not immediately as for (i), we obtain intermediate cases of waiting boundary conditions with 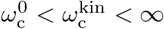. In mammalian cells, such regulating mechanisms could be provided by the kinesin MCAK, which is localized at the centromere during metaphase [41] and has been reported to increase the catastrophe rate of MTs roughly 7-fold [42]. Therefore, waiting boundary conditions with an increased catastrophe rate appear to be the most realistic scenario. We introduce a numerical catastrophe enhancement factor *n* ≥ 1 characterizing the increased catastrophe rate, 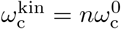. Within this general scenario reflecting boundary conditions (i) are recovered for *n* = ∞ and (ii) waiting boundary conditions with the zero force catastrophe rate for *n* = 1. We will discuss the general case (iii) in the following.

In our basic model, where MTs can exert pushing forces on kinetochores, the pushing phases where *x* > 0 can also be interpreted as a an effective waiting phase at the kinetochore with a catastrophe rate, which is effectively increased by the pushing forces. Therefore, the behavior of our basic model resembles a model with waiting boundary conditions with an increased catastrophe rate *n* > 1 at the kinetochore. MT pushing forces are not essential for bistability and oscillations and have a similar effect as an increased catastrophe rate at the kinetochore as our detailed analysis will show.

In the Fokker-Planck solution for the one-sided model, all confining boundary conditions limit the maximum MT-kinetochore distance *x*_max_ to zero, where it is positive in the basic model. When *x*_max_ is negative in the basic model (for 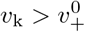, see Table 2), confining boundary conditions do not modify the basic model, since the MTs are not able to reach the fast kinetochore. For negative kinetochore velocities 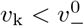, the minimum distance *x*_min_ becomes positive while *x*_max_ is zero. Then, all confining boundary conditions fix the MT tips to the kinetochore position as they do not shrink fast enough to move away from the poleward-moving kinetochore after a catastrophe resulting in 〈*x*〉 = 0 and *F*_ext_ = *γv*_k_. All in all, confinement leads to the following maximal and minimal values for the MT-kinetochore distance *x* modifying Table 2:

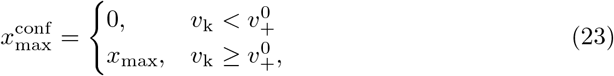

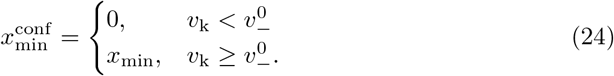

We calculate the master curves 〈*x*〉(*v*_k_) for all three types of confining boundary conditions (see Fig 11A). Because 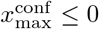 for any confining boundary condition, also 〈*x*〉 < 0, i.e., the complete master curves lie in the regime of tensile MT-kinetochore linker forces reflecting the fact that pushing forces are strictly suppressed. Therefore, the MT-kinetochore catch bond is on average under tension establishing a more firm MT-kinetochore connection during the stochastic chromosome oscillations in metaphase. Oscillations then become a tug-of-war, in which both sets of MTs only exert pulling forces onto each other.

With a waiting boundary condition at the kinetochore, the probability densities *p_±_*(*x, t*) have to be supplemented with the probability *Q*(*t*) to find a MT at the kinetochore (*x* = 0). Besides the FPEs (5) and (6) for the probability densities, we also have to solve the equation for the time evolution of *Q*(*t*):

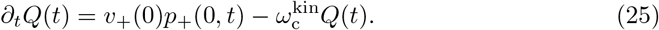

The analogous model for a free MT that grows against a rigid wall has already been solved in Refs. [31, 43]. In the stationary state, Eq (25) leads to 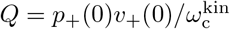. For the probability densities *p*_±_(*x*) we get the same solution as for the basic model without confinement, except for the normalization constant. The overall probability density can then be written as *p*(*x*) = *p*_+_(*x*) + *p*_−_(*x*) + *Qδ*(*x*) and has to satisfy 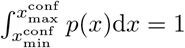.

From the overall probability density *p*(*x*) we obtain the master curves, which we show in Fig 11A for *n* = 1, 5, 20, 50, 200, ∞ and a linker stiffness of *c* = 20 pN μm^−1^. Again we can analyze the master curves for extrema to obtain constraints on linker stiffness *c* and catastrophe enhancement factor 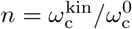 for the occurrence of bistability and oscillations. The results of this analysis are shown in Fig 11B as colored regions. It turns out that extrema in the master curve and, thus, bistability occur if the linker stiffness is sufficiently high *c* > *c*_bist_. For the zero force catastrophe rate *n* = 1 we find a high threshold value *c*_bist_ = 178 pN μm^−1^, in the limit of a reflecting boundary *n* = ∞ a very low threshold *c*_bist_ = 1.218 pN μm^−1^.

We remind that a sufficient condition for oscillations is the absence of a stable fixed point, where one of the two branches in the *v*_k_-*F*_kk_-diagram crosses *v*_k_ = 0. In contrast to the basic model, the maxima of the master curve are now located at a positive velocity for *n* > 1. Therefore, oscillations are suppressed by a fixed point 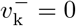 on the lower branch in the *v*_k_-*F*_kk_-diagram, which occurs if the velocity is positive in the minimum of the master curve. In general, oscillations occur if the linker stiffness is sufficiently high *c* > *c*_osc_. Again we find a high threshold value *c*_osc_ = 280 pN μm^−1^ for *n* = 1 and a low threshold *c*_osc_ = 1.237 pN μm^−1^ for a reflecting boundary condition (*n* = ∞).

**Fig 11.**
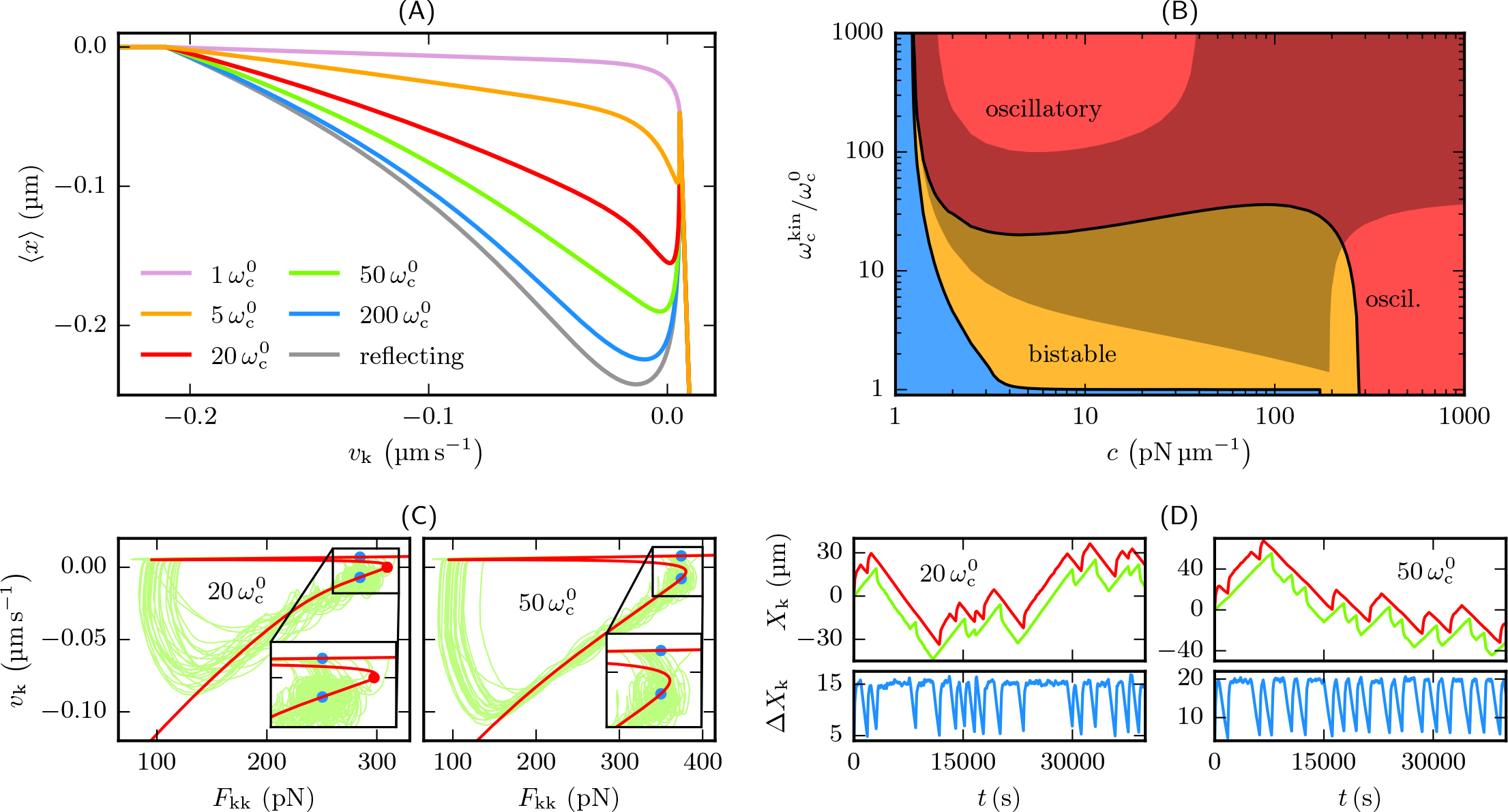
Microtubule confinement at the kinetochore. (A) Master curves of a system with a waiting boundary condition for various 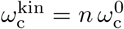 and *c* = 20 pN μm^−1^. (B) Regimes in the parameter plane of *c* and 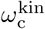 in the limit of many MTs. Outside the blue region, the master curve is bistable. In the orange region, the left branch of the master curve and, therefore, the lower branch of the *v*_k_-*F*_kk_-diagram cross *v*_k_ = 0, which leads to a fixed point suppressing oscillations (see text), whereas in the red region oscillations are possible. In stochastic simulations, kinetochores already oscillate at much smaller 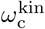 than predicted by the master curves. Additionally, a new kind of fixed point, which is depicted in (C), emerges in the shaded region. (C,D) Phase space diagrams and kinetochore trajectories from simulations of the unconfined two-sided model with *c* = 20 pN μm^−1^ and *M* = 100. The blue dots mark the new kind of fixed point, where the leading kinetochore in the lower branch moves with the same velocity as the trailing kinetochore in the upper branch. Then the inter-kinetochore distance remains constant, while the center of mass moves with a constant velocity as in (D) for 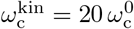 at *t* ≈ 25 000 s. In the presence of PEFs, these fixed points are absent and the shaded region in (B) does not apply.

For *n* < 10 the threshold values remain high. Moreover, at such high linker stiffnesses and for for small *n*, the simulations of the two-sided model do not show the expected behavior. For *n* = 1 and high linker stiffnesses in the oscillatory regime the kinetochore trajectories do not exhibit regular oscillations. Naively, one could argue that kinetochore oscillations are suppressed due to the lack of a pushing force and can be restored by additional PEFs. However, this is not the case, since, as stated above, PEFs do not affect the master curve that determines the regime of kinetochore motion. One reason for the absence of oscillations is that, for the zero force catastrophe rate (*n* = 1) the waiting time 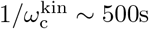 (see Table 1) at the kinetochore is large compared to the typical oscillation periods, which are in the range of 100 − 200s.

Figure 11B also shows that oscillations require increased catastrophe rates with *n* ≳ 20 over a wide range of linker stiffnesses from *c* = 10 pN μm^−1^ to *c* = 200 pN μm^−1^. For *n* > 1, at the boundary between bistable and oscillatory regime in Fig 11B, a fixed point 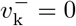 on the lower branch of the *v*_k_-*F*_kk_ phase space diagrams appears, which can suppress oscillations. This fixed point is, however, less relevant because the kinetochores will only occasionally pass the lower branch simultaneously, which is necessary to reach this fixed point. Furthermore, this fixed point is located near the right turning point *F*_max_ so that the kinetochores can easily leave the fixed point by a stochastic fluctuation (as in Fig 6). For these two reasons, in stochastic simulations, oscillations already occur for *n* ≳ 5, that is at a much lower *n* than the deterministically predicted *n* ≳ 20, but not for *n* = 1, i.e., in the absence of a catastrophe promoting mechanism.

The fixed point analysis of the *v*_k_-*F*_kk_ phase space diagrams reveals that also a new type of fixed point corresponding to a non-oscillatory motion emerges for *n* ≲ 100 in the shaded regions in Fig 11B. In this new type of fixed point, the leading P-moving kinetochore in the lower branch of the master curve has the same velocity as the trailing AP-moving kinetochore in the upper branch (see Fig 11C) so that 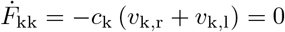, and the inter-kinetochore distance remains constant, while the center of mass moves with a constant velocity (see Fig 11D). In the presence of PEFs, however, this new type of fixed point does not survive because for the P-moving kinetochore the AP-directed PEFs increase, whereas they decrease for an AP-moving kinetochore. Then the upper blue dot in Fig 11C moves to the left, while the lower blue point moves to the right such that this new type of fixed point is unstable in the presence of PEFs. Therefore, in the entire shaded region in Fig 11B PEFs are essential to re-establish oscillations.

We conclude that both the linker stiffness *c >* 10 pN μm^−1^ and the catastrophe rate 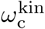 at the kinetochore (*n* ≳ 20 or *n* ≳ 5 in the presence of stochastic fluctuations) have to be sufficiently large to obtain bistability and oscillations. Because additional catastrophe promoting proteins are necessary to increase the catastrophe rate at the kinetochore, the lowest values of *n*, which still enable oscillations, might be advantageous in the cellular system. We note that poleward flux with a sufficiently large flow velocity can establish additional fixed points on the upper branch of the phase space diagrams, which suppress oscillations as in the basic model.

Moreover, the linker stiffness has to be sufficiently high to give linker extensions compatible with experimental results. An important part of the MT-kinetochore linkage is Ndc80, which is a rod-like fibril of total length around 60 nm [44, 45] consisting of two coiled-coil regions with a flexible hinge that can adopt bending angles up to 120° with a broad distribution [45]. This bending corresponds to linker length changes of *x* 50 ~ nm. Moreover, fluorescent labeling showed total intra-kinetochore stretches around 100 nm [46] or 50 nm [12]. Therefore, we regard linker extensions *x* ≲ 100 nm as realistic values. For large *n* ≫ 20 only a small linker stiffness is necessary to enable oscillations. At the small threshold stiffness, the average linker length |〈*x*〉| is typically 1 μm in this regime. Increasing the linker stiffness leads to a decreasing linker length |〈*x*〉|. We conclude that, for *n* ≫ 20, experimental observations of linker extensions |*x*| ≲ 100 nm put a stronger constraint on linker stiffness than the experimental observations of oscillations. Linker stiffnesses significantly above 5 pN μm^−1^ and, thus, far above *c*_osc_ are necessary to obtain a realistic linker length.

For *n* ~ 10 − 20, which is compatible with the experimental result *n* ~ 7 for the catastrophe promoter MCAK [42], and a linker stiffness *c* = 20 pN μm^−1^, the increased catastrophe rate at the kinetochore leads to a realistic behavior with linker extensions *x* ~ 100 nm, which are also compatible with the experimental results [12, 44–46] (see Fig 11A). This parameter regime is within the shaded regions in Fig 11B and PEFs are necessary to establish oscillations. The linker extension is independent of PEFs.

For an increased catastrophe rate around *n* ~ 10 − 20 and a linker stiffness *c* = 20 pN μm^−1^, the more realistic model with waiting boundary conditions at the kinetochore exhibits a similar behavior as our basic model because pushing phases where *x* > 0 in the basic model have a similar duration as waiting times at the kinetochore in the more realistic model.

## 9 Discussion

We provided an analytical mean-field solution of the one-sided spindle model introduced by Banigan *et al.* [24], which becomes exact in the limit of large MT numbers. The mean-field solution is based on the calculation of the mean linker extension 〈*x*〉 as a function of a constant kinetochore velocity *v*_k_ (the master curve). Together with the equation of motion of the kinetochore we obtained the force-velocity relation of the one-sided model from the master curve.

Interpreting the bistable force-velocity relation as phase space diagram, we were able to deduce idealized kinetochore oscillations, whose periods conform with experimental results [11]. For a MT-kinetochore linker stiffness *c* = 20 pN μm^−1^ and 20–25 MTs per kinetochore, we get periods of 206–258 s and 103–129 s for kinetochore and breathing oscillations, respectively. Bistability of the force-velocity relation is a necessary (but not sufficient) condition for oscillations. Our approach reproduced the frequency doubling of breathing compared to single kinetochore oscillations, observed in the experiment [11]. Both in the model and in the experiment this doubling originates from the different velocities of AP- and P-moving kinetochores, which ensure that a P-to-AP switch always follows an AP-to-P switch. In the model the velocity difference is, however, much larger. As a consequence, in our model with 20–25 MTs an AP-to-P switch follows 96–119 s after a P-to-AP switch of the sister kinetochore, which is 93 % of a breathing period, whereas in PtK2 cells a mean interval of merely 6 s has been measured [12]. In our model, different AP- and P-velocities are based on the fact that the MT shrinkage is much faster than growth. The model parameters for MT dynamics were taken from experimental measurements with yeast kinetochores [2], which, however, are distinct from metazoan kinetochores in two main points: firstly, they can only attach to one MT [47]; secondly, the Ndc80 fibrils are connected to MTs via ring-like Dam1 complexes, which do not appear in metazoan cells [48]. Therefore, it is doubtful whether yeast MT parameters can be used for modeling metazoan spindles and the model trajectories may be fitted to the experimental ones by adjusting these parameters.

In experiments with HeLa cells Jaqaman *et al.* [49] observed an increase of oscillation amplitudes and periods when they weakened the cohesin bond. In our model, a smaller cohesin stiffness *c*_k_ has the same two effects as the inter-kinetochore distance has to be larger to reach the turning points *F*_min_ and *F*_max_ of the hysteresis loop, and the phase space velocity 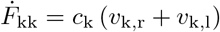 and, therefore, the frequencies are proportional to *c*_k_.

Our analytical approach also allowed us to go beyond the results of Ref. [24] and quantify constraints on the linker stiffness *c* and the MT number for occurrence of bistability in the one-sided model and for the occurrence of oscillations in the full model. We found that bistability requires linker stiffnesses above *c*_bist_ ≃ 8 pN μm^−1^. Bistability is, however, not sufficient for oscillations. Our phase space interpretation showed that bistability only leads to directional instability if the two branches of the force-velocity relation are also separated by the zero velocity line. This condition quantifies the oscillatory regime in the parameter plane of *c* and *M*. We predict that oscillations should only be observable if the MT-kinetochore linker stiffness is above *c*_osc_ ≃ 16 pN μm^−1^. Our model can thus provide additional information on the MT-kinetochore linkers whose molecular nature is unknown up to now. Several Ndc80 fibrils, which cooperatively bind to the MT, are an important part of the MT-kinetochore link and the stiffness of this Ndc80 link has been determined recently using optical trap measurements [50]. These experiments found stiffnesses above ~20 pN μm^−1^, which are compatible with our bounds. Moreover, they found a stiffening of the link under force, which could be included in our model in future work.

The derivation of the lower bound for the stiffness for the occurrence of oscillations is based on the occurrence of a new zero AP-velocity fixed point in the force-velocity diagram of the kinetochores, which suppresses oscillations upon decreasing the stiffness. Also the influence of poleward flux to the system could be analyzed by a fixed point analysis of the force-velocity diagram. Since poleward MT flux shifts the force-velocity towards smaller AP-velocities of the kinetochore, the upper branch may cross zero velocity establishing again a zero velocity fixed point suppressing oscillations. This explains why high flux velocities suppress directional instability and rationalizes the correlation between kinetochore oscillations and poleward flux observed in several cells (Table 3).

Also experimental results in Ref. [51] on the effects of phosphorylation of Hec1 onto kinetochore dynamics can be rationalized by our force-velocity diagram of the kinetochores. A lack of phosphorylation of Hec1 damps oscillations and leads to firm attachment of kinetochores to microtubules [51]. This could impede depolymerization leading to an upward shift of the lower branch of the force-velocity curve towards higher velocities, which can lead to a loss of bistability.

Furthermore, we added linearly distributed PEFs, which depend on the absolute kinetochore positions. Their main effect is a phase shift between the sister kinetochores in their phase space trajectories, which leads to regularly alternating kinetochore oscillations and, finally, forces the kinetochores to stay near the spindle equator. Consistently, experimental results show that a proper formation of the metaphase plate is not assured when PEFs are suppressed [40]. Since the PEFs do not affect the master curves and phase space diagrams, deterministically, they never completely suppress oscillations but only reduce their amplitude and increase their frequency. In stochastic simulations, the kinetochore oscillations are more fluctuative in the presence of PEFs, but are only completely suppressed for rather strong PEFs. A similar observation was made in the model of Civelekoglu-Scholey *et al.* [23].

Finally, we lifted the assumption that MTs are able to apply pushing forces on the kinetochores because experiments show that MTs only exert tensile forces [7, 25]. Therefore, we confined MT growth at the kinetochore by catastrophe-triggering boundary conditions. The catastrophe rate for a MT at the kinetochore 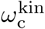 can, in principle, range from the force-free MT catastrophe rate 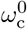, which is realistic in the absence of any catastrophe promoting proteins up to infinity if a catastrophe is immediately triggered. In the presence of the centromere-associated regulating protein MCAK increased catastrophe rates 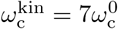 are expected [42]. We found that *both* the linker stiffness *c and* the catastrophe rate 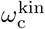 at the kinetochore have to be sufficiently large to obtain bistability and oscillations. We find, in particular, that the force-free MT catastrophe rate is not sufficient to lead to oscillations, which shows that catastrophe-promoting proteins are essential to induce oscillations. In the presence of PEFs, oscillations can be recovered also for relatively small catastrophe rates: For 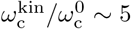, we found no oscillations in the absence of PEFs; for 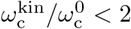 we found no oscillations at all. Moreover, the linker stiffness has to be sufficiently high to give linker extensions below 100 nm compatible with experimental results [12, 44–46]. For 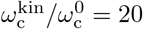 and a linker stiffness of *c* = 20 pN μm^−1^, we found realistic behavior. Our results can explain experimental observations in Ref. [52], where PtK2-cells were observed under depletion of centromeric MCAK, which decreases 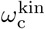. Then, in accordance to our results (see Fig 11CD), the turning point *F*_max_ of the hysteresis loop decreases. As a result the oscillation frequency increases and the mean centromere stretch decreases, while the “motility rates”, i.e., the velocities do not change.

Kinetochore motion in the non-oscillatory regime can be described as fluctuations around a fixed point with constant inter-kinetochore distance. This is exactly the behavior of peripheral kinetochores in PtK1 cells [11], while the central kinetochores do exhibit directional instability. Civelekoglu-Scholey *et al.* [23] explained this dichotomy with different distributions of polar ejection forces in the center and the periphery of the metaphase plate. This explanation is also consistent with our stochastic simulations. Our results suggest, however, also differences in linker stiffness or catastrophe promotion as possible reasons for the dichotomy. For instance, less Ndc80 complexes could participate in peripheral MT-kinetochore links resulting in reduced linker stiffness and non-oscillatory behavior. Also a non-uniform MCAK distribution that decreases radially towards the periphery of the metaphase plate could reduce 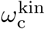 and suppress oscillations of peripheral kinetochores. Differences in poleward flux might be another possible explanation for the dichotomy according to our results. However, Camereon *et al.* [53] observed that the flux velocities in PtK1 cells do not depend on the chromosome to which a microtubule is attached.

In conclusion, the minimal model can rationalize a number of experimental observations. Particularly interesting are constraints on the MT-kinetochore linker stiffness that are compatible with recent optical trap measurements [50]. The predicted responses to the most relevant parameter changes are summarized in Table 4 and suggest further systematic perturbation experiments, for example, by promoting catastrophes at the kinetochore.

**Table 4.**
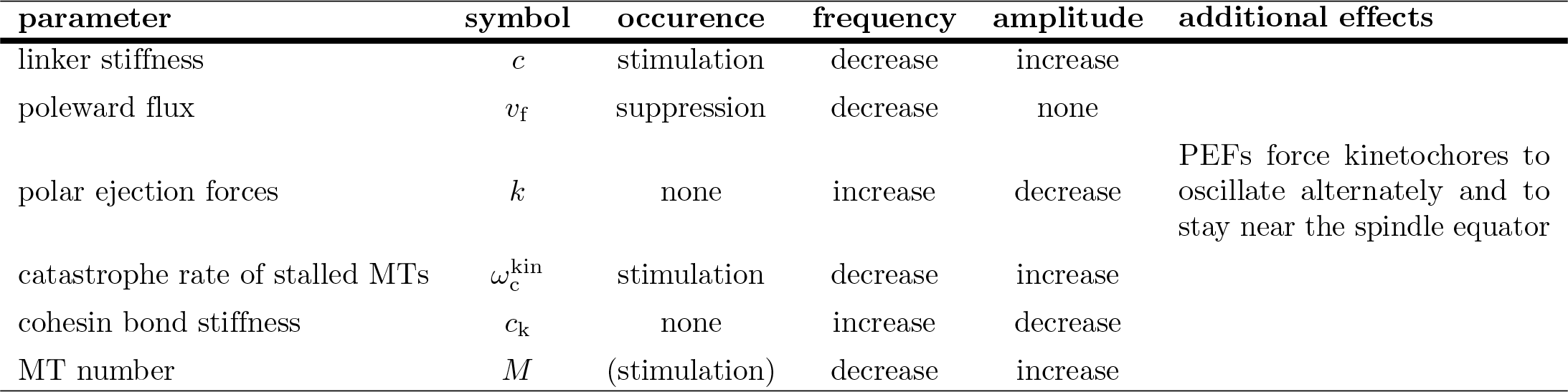
Summary. Effect of an increase of the parameter in the first column on occurrence, frequency, and amplitude of kinetochore oscillations.

| parameter | symbol | occurrence | frequency | amplitude | additional effects |
| --- | --- | --- | --- | --- | --- |
| linker stiffness | $c$ | stimulation | decrease | increase | |
| poleward flux | $v_f$ | suppression | decrease | none | |
| polar ejection forces | $k$ | none | increase | decrease | PEFs force kinetochores to oscillate alternately and to stay near the spindle equator |
| catastrophe rate of stalled MTs | $\omega_c^{\text{kin}}$ | stimulation | decrease | increase | |
| cohesin bond stiffness | $c_k$ | none | increase | decrease | |
| MT number | $M$ | (stimulation) | decrease | increase | |

## Supporting information

**S1 Appendix. Supplemental derivations.** Additional material on linker length distributions and an alternative mean-field approach.

**S1 Video. Deterministic kinetochore motion in the basic model.** Illustration of kinetochore motion in the force-velocity plane (*F*_kk_-*v*_k_ phase space diagrams) and in real space. The used parameters are *c* = 20 pN μm^−1^ and *M* = 25 as depicted in Fig. 3.

**S2 Video. Stochastic kinetochore motion in simulations of the basic model.** Illustration of kinetochore motion in the *F*_kk_-*v*_k_ phase space and in real space. Simulations were perfomed with an unconfined system, *c* = 20 pN μm^−1^ and *M* = 25 as depicted in Fig. 4B.

**S3 Video. Stochastic kinetochore motion in simulations of the basic model.** Illustration of kinetochore motion in the *F*_kk_-*v*_k_ phase space and in real space. Simulations were perfomed with an unconfined system, *c* = 20 pN μm^−1^ and *M* = 500 as depicted in Fig. 4B.

**S4 Video. Deterministic kinetochore motion under the influence of PEFs.** Illustration of kinetochore motion in the *F*_kk_-*v*_k_ phase space and in real space. The used parameters are *c* = 20 pN μm^−1^, *M* = 25 and *k* = 10 pN μm^−1^ as depicted in Fig. 9.

**S5 Video. Stochastic kinetochore motion under the influence of moderate PEFs.** Illustration of kinetochore motion in the *F*_kk_-*v*_k_ phase space and in real space. Simulations were perfomed with an unconfined system, *c* = 20 pN μm^−1^, *M* = 100 and *k* = 10 pN μm^−1^ as depicted in Fig. 10.

**S6 Video. Stochastic kinetochore motion under the influence of strong PEFs.** Illustration of kinetochore motion in the *F*_kk_-*v*_k_ phase space and in real space. Simulations were perfomed with an unconfined system, *c* = 20 pN μm^−1^, *M* = 100 and *k* = 1000 pN μm^−1^ as depicted in Fig. 10.

