## Supplementary material for "Bistability and oscillations in cooperative microtubule and kinetochore dynamics in the mitotic spindle": S1 Appendix

Felix Schwietert* and Jan Kierfeld†
*Physics Department, TU Dortmund University, 44221 Dortmund, Germany*
(Dated: February 21, 2019)

In this Supplemental Material we present additional material on linker length distributions and an alternative mean-field approach.

### I. BEHAVIOR OF LINKER LENGTH DISTRIBUTIONS NEAR THE MAXIMUM AND MINIMUM LINKER EXTENSIONS

For $v_\mathrm{k} > 0$ ($v_\mathrm{k} < 0$) the linker length distributions $p_+(x,t)$ ($p_-(x,t)$) as given by Eq (12) in the main text,

$$p_\pm(x) = \frac{\pm\mathcal{N}}{v_\pm(x)} \exp\left(-\int \left(\frac{\omega_\mathrm{c}(x)}{v_+(x)} + \frac{\omega_\mathrm{r}(x)}{v_-(x)}\right)\mathrm{d}x\right), \tag{S1}$$

have singularities or peaks at $x = x_\mathrm{max}$ ($x = x_\mathrm{min}$) where $v_+(x_\mathrm{max}) = 0$ ($v_-(x_\mathrm{min}) = 0$), see Fig S1. These singularities occur because the integral

$$I(x) \equiv -\int \left(\frac{\omega_\mathrm{c}(x)}{v_+(x)} + \frac{\omega_\mathrm{r}(x)}{v_-(x)}\right)\mathrm{d}x, \tag{S2}$$

in the exponent and the prefactor $1/v_\pm(x)$ in the expression (Eq S1) diverge for $v_+(x_\mathrm{max}) = 0$ ($v_-(x_\mathrm{min}) = 0$).

To investigate the nature of these singularities or peaks in more detail, we expand around $x_\mathrm{max}$ and $x_\mathrm{min}$ to leading order, starting with $x_\mathrm{max}$. Since $v_-(x_\mathrm{max}) \neq 0$, the second term of $I(x)$ in Eq (S2) simply yields a finite factor

$$\beta(x) \equiv \exp\left(-\int \frac{\omega_\mathrm{r}(x)}{v_-(x)}\mathrm{d}x\right).$$

For the first term, we find for $x \lessapprox x_\mathrm{max}$

$$\frac{\omega_\mathrm{c}(x)}{v_+(x)} \approx \frac{\alpha_+ + 1}{x_\mathrm{max} - x} \quad \text{with}$$
$$\alpha_+ + 1 = \frac{\omega_\mathrm{c}^0 F_+}{c v_+^0}\left(\frac{v_\mathrm{k}}{v_+^0}\right)^{-1+F_+/F_\mathrm{c}} > 0$$

resulting in

$$I(x) \approx (\alpha_+ + 1)\ln(x_\mathrm{max} - x) + \ln\beta(x),$$
$$\exp(I(x)) \approx \beta(x_\mathrm{max})(x_\mathrm{max} - x)^{\alpha_+ + 1}.$$

Because $\alpha_+ + 1 > 0$ we have for $x \lessapprox x_\mathrm{max}$

$$p_-(x) \approx -\frac{\mathcal{N}}{v_-(x_\mathrm{max})}\exp(I(x)) \propto (x_\mathrm{max} - x)^{\alpha_+ + 1} \approx 0,$$

and, therefore, $p(x) \approx p_+(x)$. Analyzing the prefactor

$$\frac{1}{v_+(x)} \approx \frac{\mathrm{const}}{x_\mathrm{max} - x} > 0,$$

we finally find a power-law behavior

$$p(x) \approx p_+(x) \approx \mathcal{N}\mathrm{const}\,\beta(x_\mathrm{max})(x_\mathrm{max} - x)^{\alpha_+}$$
$$\propto (x_\mathrm{max} - x)^{\alpha_+}$$

for $x$ approaching $x_\mathrm{max}$.

In an analogous manner, we expand around $x = x_\mathrm{min}$ to leading order and find a power-law behavior

$$p(x) \approx p_-(x) \propto (x - x_\mathrm{min})^{\alpha_-} \quad \text{with}$$
$$\alpha_- + 1 = \frac{\omega_\mathrm{r}^0 F_-}{c v_-^0}\left(\frac{v_\mathrm{k}}{v_-^0}\right)^{-1+F_-/F_\mathrm{r}} > 0.$$

The resulting dependencies of the exponents $\alpha_\pm + 1 \propto 1/c$ on $c$ and $\alpha_\pm + 1 \propto (|v_\mathrm{k}/v_\pm^0|)^{-1-|F_\pm/F_{\mathrm{c,r}}|}$ on kinetochore velocities $|v_\mathrm{k}|$ are used and discussed in the main text. Since $\alpha_\pm > -1$, the probability densities are always normalizable despite the singularities at $x_\mathrm{max}$ and $x_\mathrm{min}$.

If $\alpha_\pm < 0$ (for sufficiently large $c$ or sufficiently large $|v_\mathrm{k}|$), the resulting total distributions $p(x) = p_+(x) + p_-(x)$ are peaked around $x_\mathrm{max}$ or $x_\mathrm{min}$. In the unstable regime around $v_\mathrm{k} \approx 0$, however, the linker length distribution $p(x)$ becomes broad without pronounced peaks. This behavior is visualized in Fig S1. In this regime, the kurtosis $\langle(x - \langle x\rangle)^4\rangle / \langle(x - \langle x\rangle)^2\rangle^2$, which is a measure of the sharpness of the peaks of the distribution $p(x)$ around $x_\mathrm{min}$ and $x_\mathrm{max}$, becomes minimal indicating a broad distribution $p(x)$.

### II. MEAN-FIELD THEORY FOR THE ONE-SIDED MODEL ASSUMING IDENTICAL LINKER EXTENSIONS

Here, we present an alternative but simpler mean-field approximation for the one-sided model. We assume that all linkers have *identical* extensions ($x_i \approx x$ for all $i$), i.e., all MTs have identical lengths and are in the same state (growing or shrinking). While we assume in the mean-field approach presented in the main text that all MTs

*
†

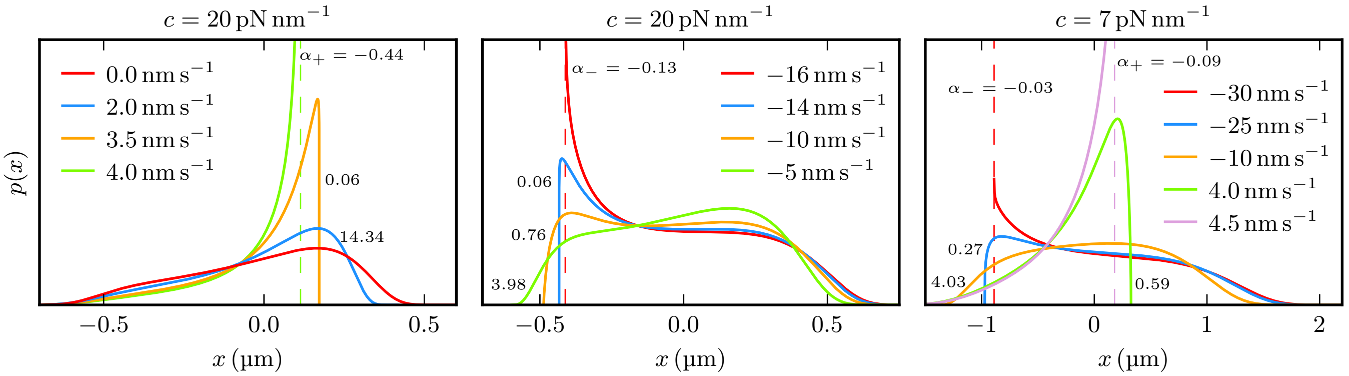


FIG. S1. Probability densities of linker extensions $p(x) = p_+(x) + p_-(x)$ due to Eq (S1), for various combinations of linker stiffnesses $c$ and kinetochore velocities $v_\mathrm{k}$. For $v_\mathrm{k} > 0$ ($v_\mathrm{k} < 0$), they show the predicted power law behavior at $x_\mathrm{max}$ ($x_\mathrm{min}$) that is determined by $\alpha_+$ ($\alpha_-$). For $c = 20\,\mathrm{pN\,nm^{-1}}$, the distributions are peaked ($\alpha_\pm < 0$) if $v_\mathrm{k} > 3.54\,\mathrm{nm\,s^{-1}}$ ($v_\mathrm{k} < 14.6\,\mathrm{nm\,s^{-1}}$). Thus, the mean linker extension stays near $x_\mathrm{max}$ ($x_\mathrm{min}$) also in the vicinity of $v_\mathrm{k} = 0$ so that it has to have a positive slope to get from $x_\mathrm{min}$ to $x_\mathrm{max}$ during the evolution from negative to positive velocities, resulting in a bistability. For $c = 7\,\mathrm{pN\,nm^{-1}}$, which lies below the regime boundary $c_\mathrm{bist}$, the distributions themselves as well as the unpeaked region ($\alpha_\pm > 0$, $-29.3\,\mathrm{nm\,s^{-1}} < v_\mathrm{k} < 4.41\,\mathrm{nm\,s^{-1}}$) are broader so that the mean linker extension evolves monotonically from $x_\mathrm{min}$ to $x_\mathrm{max}$ when $v_\mathrm{k}$ is increased and crosses $v_\mathrm{k} = 0$.

approximately *decouple* as soon as kinetochore velocity fluctuations are neglected ($v_\mathrm{k} = \mathrm{const}$), we assume here a *strong coupling* between MTs. Accordingly the compound linker distribution does no longer factorize but can still be described by a single function $p_\pm(x,t)$, which is the probability to find *all* MTs in the growing (+) or shrinking (−) state with a MT-kinetochore linker extension $x$.

While this approximation appears much more restrictive regarding the MT length (and thus the linker extension) distribution, it allows us to include stochastic fluctuations of the kinetochore velocity, which we neglected in the mean-field approach in the main text. Here, the kinetochore velocity is a stochastic variable, depending on the stochastic (but identical) linker extension $x$:

$$v_\mathrm{k} = \frac{1}{\gamma}\left(F_\mathrm{ext} + cMx\right). \tag{S3}$$

The Fokker-Planck-equations for the probability densities $p_\pm(x,t)$ are the same as for the $v_\mathrm{k} = \mathrm{const.}$ approximation (Eqs (5) and (6) in the main text), but with a different relative velocity

$$\begin{aligned} v_\pm(x) &= v_{\mathrm{m}\pm}(x) - v_\mathrm{k} \\ &= v_\pm^0 \exp\left(-\frac{cx}{F_\pm}\right) - \frac{1}{\gamma}\left(F_\mathrm{ext} + cMx\right). \end{aligned} \tag{S4}$$

The maximum and minimum MT-kinetochore distances $x_\mathrm{max/min}$ are reached if the relative velocity vanishes, $v_\pm(x_\mathrm{max/min}) = 0$, and Eq (S3) is fulfilled. Then both the kinetochore and the MT-tips move with the same velocity $\tilde{v}_\pm$ given by Eq (9) in the main text:

$$\tilde{v}_\pm \equiv \frac{MF_\pm}{\gamma} W\left(\frac{\gamma v_\pm^0}{MF_\pm}\exp\left(\frac{F_\mathrm{ext}}{MF_\pm}\right)\right). \tag{S5}$$

The corresponding MT-kinetochore distances are

$$\begin{aligned} x_\mathrm{max/min} &= \left(F_\pm/c\right)\ln\left(v_\pm^0/v_\mathrm{k}\right) \\ &= -\frac{F_\pm}{c}\left(\frac{F_\mathrm{ext}}{MF_\pm} - W\left(\frac{\gamma v_\pm^0}{MF_\pm}\mathrm{e}^{F_\mathrm{ext}/MF_\pm}\right)\right), \end{aligned}$$

which agrees with Tab. 2 in the main text, if Eq (S5) is used to eliminate $v_\mathrm{k}$ in favor of $F_\mathrm{ext}$.

With the new relative velocities $v_\pm(x)$ from Eq (S4) we finally obtain a solution for the probability densities $p_\pm(x,t)$ analogous to Eq (12) in the main text. This solution can be used to calculate the mean linker extensions $\langle x\rangle$ and the mean kinetochore velocity

$$\langle v_\mathrm{k}\rangle = \frac{1}{\gamma}\left(F_\mathrm{ext} + cM\langle x\rangle\right) \tag{S6}$$

as a function of the external force $F_\mathrm{ext}$. This type of mean-field approach will never result in a bistable force-velocity relation as we always obtain a unique mean linker extension $\langle x\rangle$ and, according to Eq (S6), a unique mean kinetochore velocity as a function of the force $F_\mathrm{ext}$. In order to map out bistability, it is technically advantageous to consider $\langle x\rangle$ and, thus, $F_\mathrm{ext}$ as a function of the kinetochore velocity $v_k$ as in the mean-field approach in the main text. Then a bistable force-velocity relation can emerge as a result of a non-monotonous (but unique) mean linker extension $\langle x\rangle$ as a function of $v_k$. Nevertheless, we find hints to a bistable behaviour also in the mean-field theory with identical linker extensions: The probability density $p(x) = p_+(x) + p_-(x)$ becomes bimodal around $F_\mathrm{ext} = 0$ which corresponds to bistable temporal switching of the whole ensemble between two linker extensions and, thus, two kinetochore velocities $v_k$.

In the present approach we always assume identical linker extensions *and* identical MT states (all MTs

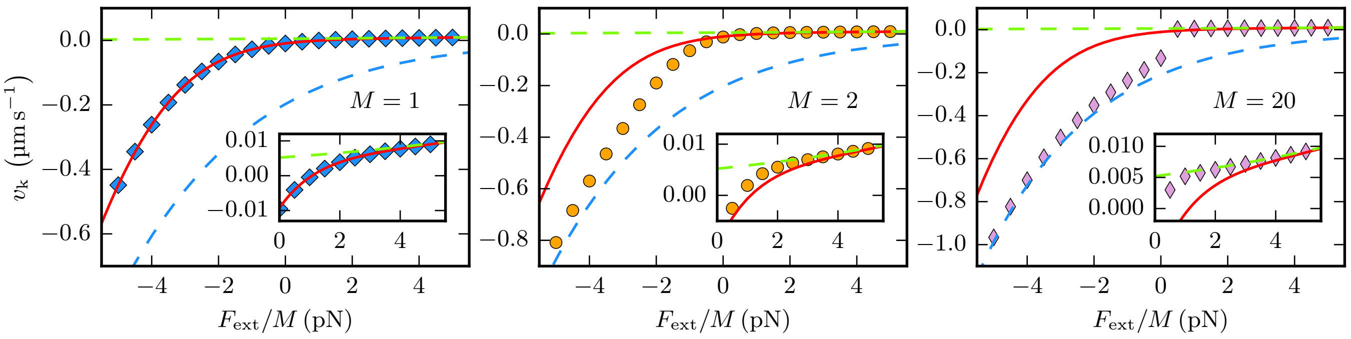


FIG. S2. Mean kinetochore velocities resulting from the approximation of identical linker extensions (red lines) compared to simulation results (markers). For a one-sided system with a single MT ($M = 1$) the alternative mean-field approach is exact by definition and gives correct mean velocities. If $M > 1$, even the simple assumption of exclusively shrinking / growing MTs [$\tilde{v}_\pm$ from Eq (S5)] gives a better approximation (green and blue dashed lines). In contrast to Fig 2, where we aimed to detect the two bistable states separately in each simulation, we average the simulations over a long period, to measure the total mean velocity.

growing or all MTs shrinking), while bistability in the $v_{\mathrm{k}} = \mathrm{const}$ mean-field approach in the main text is the result of a very broad and heterogeneous stationary linker extension distribution, where parts of the MT population switch from shrinking to growing if the velocity is increased in the bistable region around $v_{\mathrm{k}} \approx 0$. In this bistable region, not all MTs are in the same state anymore, and the assumption of identical linker extensions and states becomes invalid.

By definition, the mean-field theory with identical linker extensions is exact for a system with a single MT ($M = 1$), as can be seen in Fig S2. For an ensemble of MTs – even for the next smallest number $M = 2$ – the approach fails, however, to provide a good approximation of the mean kinetochore velocity. Then even the simple assumption of exclusively shrinking or growing MTs, which results in $v_k = \tilde{v}_\pm$, see Eq (S5), gives a better approximation in the large force regimes. For $M = 20$, which is the relevant case for mammalian cells, the mean-field theory with identical linker extensions differs strongly from the simulation results. We conclude that the $v_{\mathrm{k}} = \mathrm{const}$ mean-field approach described in the main text is superior for analyzing bistability and oscillations in the spindle model.
